# Changes in seam number and location induce holes within microtubules assembled from porcine brain tubulin and in *Xenopus* egg cytoplasmic extracts

**DOI:** 10.1101/2021.07.14.452321

**Authors:** Charlotte Guyomar, Clément Bousquet, Siou Ku, John Heumann, Gabriel Guilloux, Natacha Gaillard, Claire Heichette, Laurence Duchesne, Michel O. Steinmetz, Romain Gibeaux, Denis Chrétien

## Abstract

Microtubules are tubes of about 25 nm in diameter that are critically involved in a variety of cellular functions including motility, compartmentalization, and division. They are considered as pseudo-helical polymers whose constituent αβ-tubulin heterodimers share lateral homotypic interactions, except at one unique region called the seam. Here, we used a segmented sub-tomogram averaging strategy to reassess this paradigm and analyze the organization of the αβ-tubulin heterodimers in microtubules assembled from purified porcine brain tubulin in the presence of GTP and GMPCPP, and in *Xenopus* egg cytoplasmic extracts. We find that in all conditions, microtubules incorporate variable protofilament and/or tubulin subunit helical-start numbers, as well as variable numbers of seams. Strikingly, the seam number and location vary along individual microtubules, generating holes of one to a few subunits in size within their lattices. Together, our results reveal that the formation of mixed and discontinuous microtubule lattices is an intrinsic property of tubulin that requires the formation of unique lateral interactions without longitudinal ones. They further suggest that microtubule assembly is tightly regulated in a cytoplasmic environment.

## Introduction

The organization of the αβ-tubulin heterodimer within microtubules was originally inferred from the analysis of transmission electron microscopy images of negatively stained axonemal doublets (Amos & Klug, 1974). It was proposed that the tubulin subunits engage heterotypic lateral interactions (α-β, β-α) in the complete 13 protofilaments A-microtubule, and homotypic ones (α-α, β-β) in the incomplete 10 protofilaments B-microtubule, giving rise to the concept of the A and B lattices (Figure 1A-B). However, using kinesin-motor domains that bind uniquely to β-tubulin (Figure 1C), it was shown later that in both the A and B microtubules of the doublet, tubulin heterodimers engage homotypic interactions of the B type (Song & Mandelkow, 1995), which is also the case in microtubules assembled *in vitro* from purified tubulin (Crepeau et al., 1978; Song & Mandelkow, 1993). Noticeably, for geometrical reasons (McEwen & Edelstein, 1980; Wade & Chrétien, 1993), microtubules organized with 13 protofilaments and 3-start lateral helices should contain at least one ‘seam’ of the A-type (Figure 1B), which corresponds to our current view of microtubule lattice organization.

**Figure 1.**
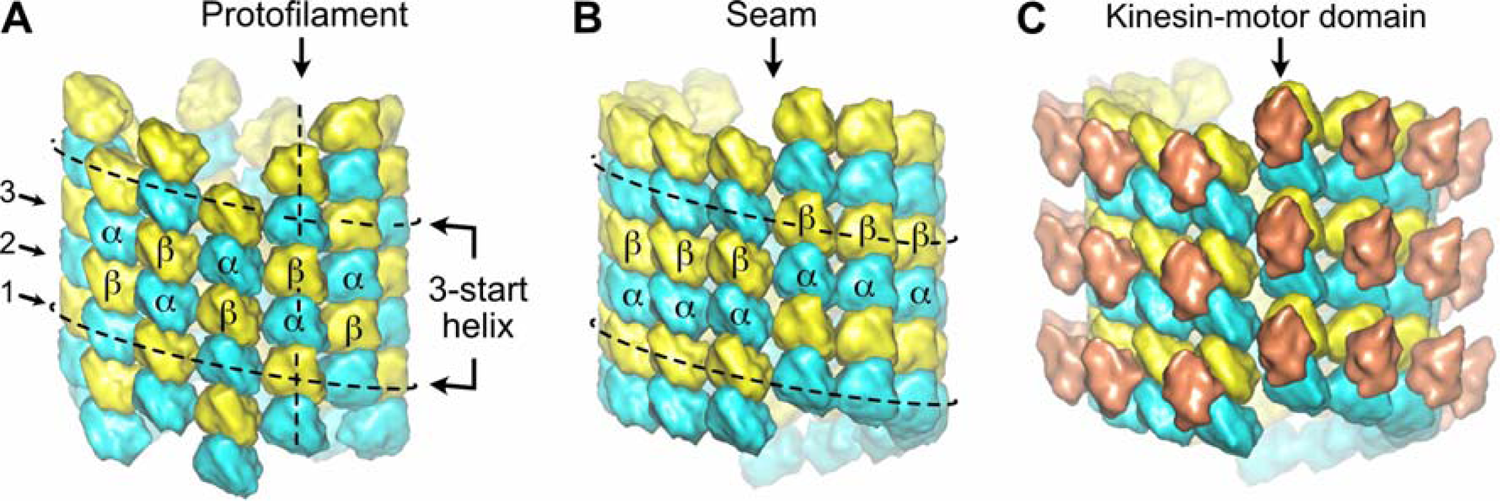
Organization of tubulin within microtubules. The αβ-tubulin heterodimers (α in cyan, β in yellow) alternate head-to-tail along protofilaments, 13 of which associate laterally to form the microtubule wall. **(A)** In the A-type lattice, the lateral contacts are made between heterotypic subunits (α-β, β-α) along the 3-start helices. **(B)** In the B-type lattice, the lateral contacts are made between homotypic subunits (α-α, β-β), except at one unique region of the A-type called the seam. **(C)** Decoration of microtubules with kinesin-motor domains (orange) that bind to β-tubulin highlights the organization of the tubulin heterodimer within microtubules.

Multiple seams were first visualized by freeze-etching and rotary shadowing of microtubules assembled *in vitro* (Kikkawa et al., 1994). Using the same approach on cells treated with detergent to remove the membrane and decorate the microtubules with kinesin-motor domains, the authors provided the first evidence of a preferred B-lattice type organization *in cellulo*, and could visualize unique seams in cytoplasmic microtubules. But due to the limitation of the method and the small number of microtubules observed, they did not exclude the possibility of several seams *in cellulo*. Since then, several studies have revealed the presence of multiple seams in microtubules assembled *in vitro,* noticeably in the presence of the stabilizing drug Taxol (Debs et al., 2020; des Georges et al., 2008; Howes et al., 2017; Sosa et al., 1997). The predominance of B-type lateral contacts *in cellulo* was confirmed by cryo-electron tomography after detergent removal of the membrane and decoration with kinesin-motor domains, but with no detailed statistics (McIntosh et al., 2009). Therefore, it turns out that our knowledge on the organization of αβ-tubulin heterodimers within microtubules assembled *in vitro* in the absence of drug and *in cellulo* remains limited.

To gain a deeper understanding of microtubule lattice organization *in vitro* and in a cytoplasmic environment, we analyzed microtubules assembled from purified porcine brain tubulin in the presence of GTP, the slowly hydrolysable analogue GMPCPP, and in *Xenopus* egg cytoplasmic extracts. Microtubules were decorated with kinesin-motor domains and their binding pattern was analyzed using cryo-electron tomography followed by sub-tomogram averaging (STA). To this end, we specifically developed a segmented sub-tomogram averaging (SSTA) strategy, which allowed us to investigate the structural heterogeneity of individual microtubules. We find that in all conditions the seam number and location vary within individual microtubules, leaving holes of one to a few subunits in size within their wall. Microtubules assembled in a cytoplasmic environment are more regular, suggesting a tightly regulated process. Moreover, the formation of discontinuous mixed AB-lattices implies that tubulin can engage unique lateral interactions without longitudinal ones at the growing tip, a process that accounts for the formation of holes within their wall during polymerization.

## Results

Microtubules were self-assembled *in vitro* from purified porcine brain tubulin in the presence of 1 mM GTP (Figure 2 - figure supplement 1A) and kinesin-motor domains were added at the polymerization plateau right before vitrification of the specimen grids into liquid ethane (Figure 2 - figure supplement 1B). Cryo-electron tomograms were acquired preferentially using a dual-axis strategy (Guesdon et al., 2013) so that all microtubules could be analyzed independently of their orientation with respect to the tilt axes (Figure 2 - figure supplement 1C, Video 1). The low magnifications used, between 25 000 X and 29 000 X, allowed us to record long stretches of the microtubules, ∼1 to 2 µm in length, to optimize the sub-tomographic averaging strategy along individual fibers.

**Figure 2.**
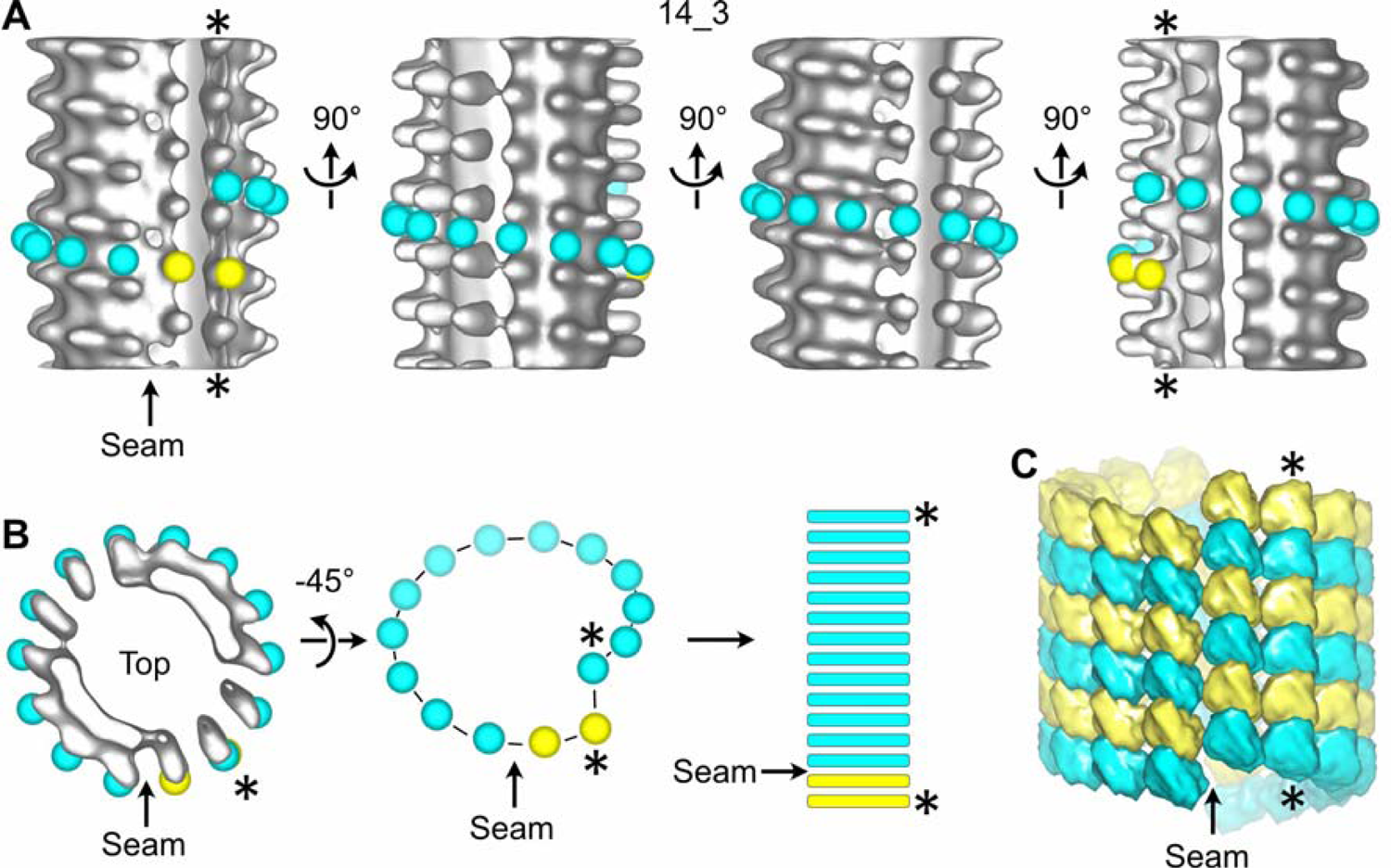
Sub-tomogram averaging of a 14_3 microtubule with a unique seam. **(A)** Sub-tomogram average of a 1390.4 nm long 14_3 microtubule assembled *in vitro* from purified tubulin and decorated with kinesin-motor domains (Figure 2 - figure supplement 3A: MT3, Video 2). The panel displays 4 views turned by 90° with respect to the longitudinal axis of the microtubule. Yellow spheres have been placed onto the kinesin-motor domain densities and cyan spheres in between them. They follow the left-handed, three-start helix of the microtubule lattice. The seam shows up as a change in color from yellow to cyan. **(B)** Symbolic representation of the microtubule lattice. The top view of the microtubule in (A) is turned by 45° around the X-axis and the density is masked to reveal the organization of the tubulin subunits in one turn of the three-start helix. The helix is unrolled and represented as longitudinal bars that correspond to the organization of the αβ-subunits in microtubule segments. **(C)** 3D model of the underlying tubulin dimer lattice. The stars (*) indicate the same protofilament in (A-C).

### The number and location of seams vary within individual microtubules assembled from purified tubulin

We first processed entire microtubules present in the tomograms using a sub-tomogram averaging approach that retrieves small sub-volumes of ∼50 nm^3^ in size at every kinesin-motor domain position (Zabeo et al., 2018) (i.e., every ∼8 nm; Figure 2 - figure supplement 2A). The resulting 3D volumes clearly revealed the protofilament number and the organization of the kinesin-motor domains around the microtubule lattice (Figure 2A, Video 2), and hence the underlying organization of their constituent tubulin dimers (Figure 2B-C). In agreement with previous studies performed on Taxol-stabilized microtubules (Debs et al., 2020; des Georges et al., 2008; Howes et al., 2017; Kikkawa et al., 1994; Sosa et al., 1997), we found that microtubules assembled *in vitro* from purified tubulin in the presence of GTP contained one to several A-lattice seams (Figure 3). However, we could frequently observe protofilaments with a much thinner appearance where the kinesin-motor domain periodicity was partly or completely lost (Figure 3A, Video 3). We hypothesized that the appearance of such aberrant protofilaments resulted from the averaging of regions containing kinesin-motor domain densities with regions falling in between. To explore this idea, we used SSTA to reconstruct short regions along individual microtubules (Figure 3B-C, Figure 3 - figure supplement 2B). Using this approach, we could identify regions where the seam number and/or location varied within individual microtubules. In the example shown in Figure 3B, the segment S1 contains 5 seams while S3 and S4 contain 3 seams. S2 still displays two aberrant protofilaments indicating that the change in seam number occurred in this region. To confirm this hypothesis, we extracted the corresponding region in the raw tomogram that was further filtered by thresholding intensities in Fourier space to increase the signal-to-noise ratio (Figure 4A). Comparison between the kinesin-motor domain patterns in the sub-tomogram averages of segments S1 to S3 with the filtered S2 region confirmed that this latter constitutes a transition zone where the seam number changes from 5 to 3. Line plots along the registered protofilaments (Figure 4B) shows that the kinesin-motor domain periodicity becomes out of phase after the transition in the aberrant protofilaments, implying an offset of at least one monomer (or an odd number) before and after the transition, and hence the presence of holes within the microtubule lattice (Figure 4C). Analysis of 24 microtubules taken on 4 tomograms, representing 195 segments of ∼160 nm length (i.e., 2664 lateral interactions), revealed an average lattice type transition frequency of 3.7 µm^-1^ (Supplementary Table 1), but with a high heterogeneity. Some microtubules showed no or little lattice type transitions (e.g., MT3 and MT4, Figure 3 - figure supplementary 3A; MT16, MT21 and MT23, Figure 3 - figure supplementary 3B), while others were heavily dislocated, with a lattice type transition frequency as high as ∼15 µm^-1^ (e.g., MT13 and MT14, Figure 3 - figure supplementary 3B).

**Figure 3.**
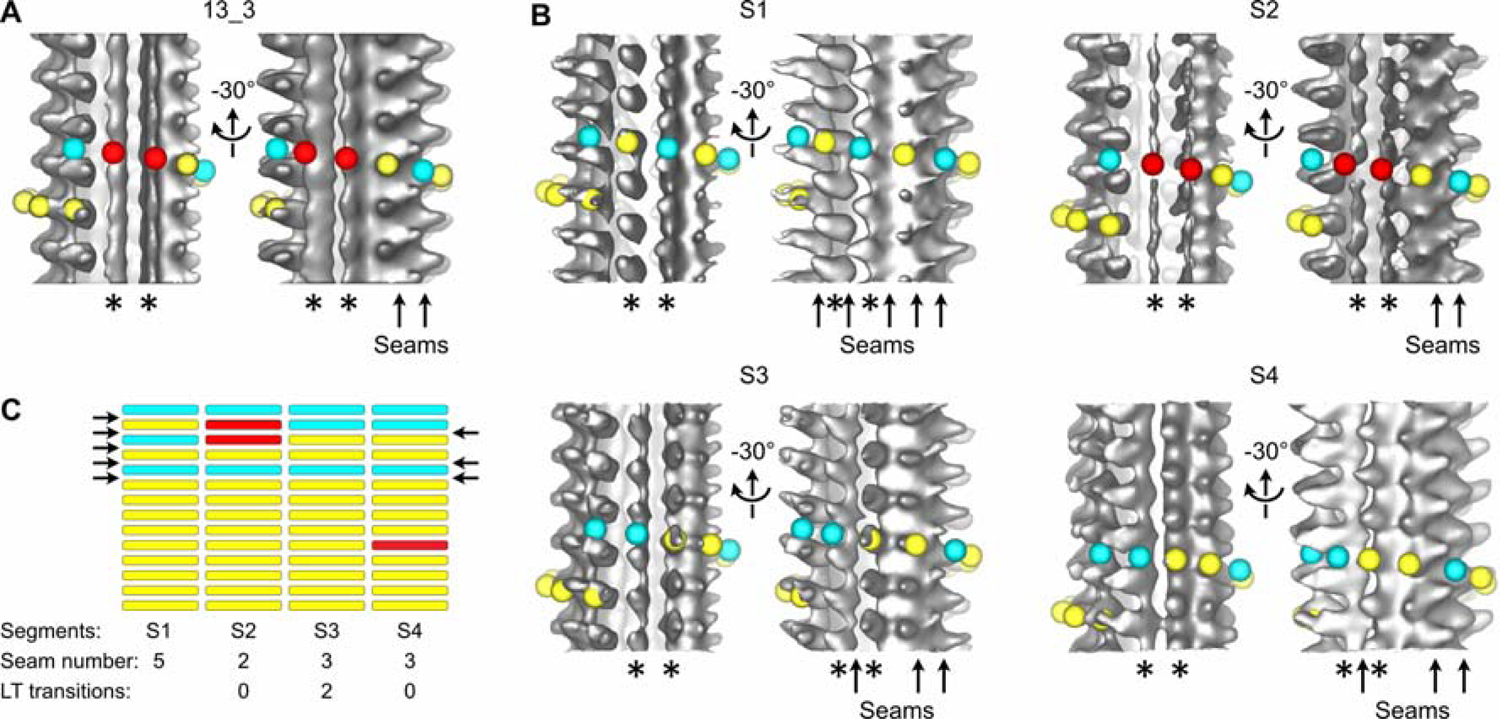
Transition in seam number within a 13_3 microtubule. **(A)** Average of a 1327.2 nm long 13_3 microtubule displaying two aberrant protofilaments (*), and two adjacent seams (arrows in the −30° view). Red spheres have been placed on top of the aberrant protofilaments. **(B)** Segmented sub-tomogram averaging of the microtubule in (A). The microtubule has been divided into 4 segments of 331.8 nm in length, and sub-tomogram averages have been calculated for each segment (S1 to S4). The two aberrant protofilaments in (A) are well resolved in S1, S3 and S4, while they still display an aberrant shape in S2. The lattice organization of these protofilaments must be offset by at least one tubulin subunit between S1 and S3. Hence, S2 constitutes a transition zone where kinesin-motor domain densities and absence of densities have been averaged. **(C)** Flat representation of the lattice organization within segments S1 to S4. S1 contains 5 seams while S3 and S4 contain 3 seams (arrows). Two lattice type (LT) transitions occur between S1 and S3, and S4 contains an aberrant protofilament (Video 3). A finer segmentation of the microtubule at 165.9 nm revealed an additional lattice type transition in this region (Figure 3 - figure supplement 3A: MT5, between segments S5 and S7).

**Figure 4.**
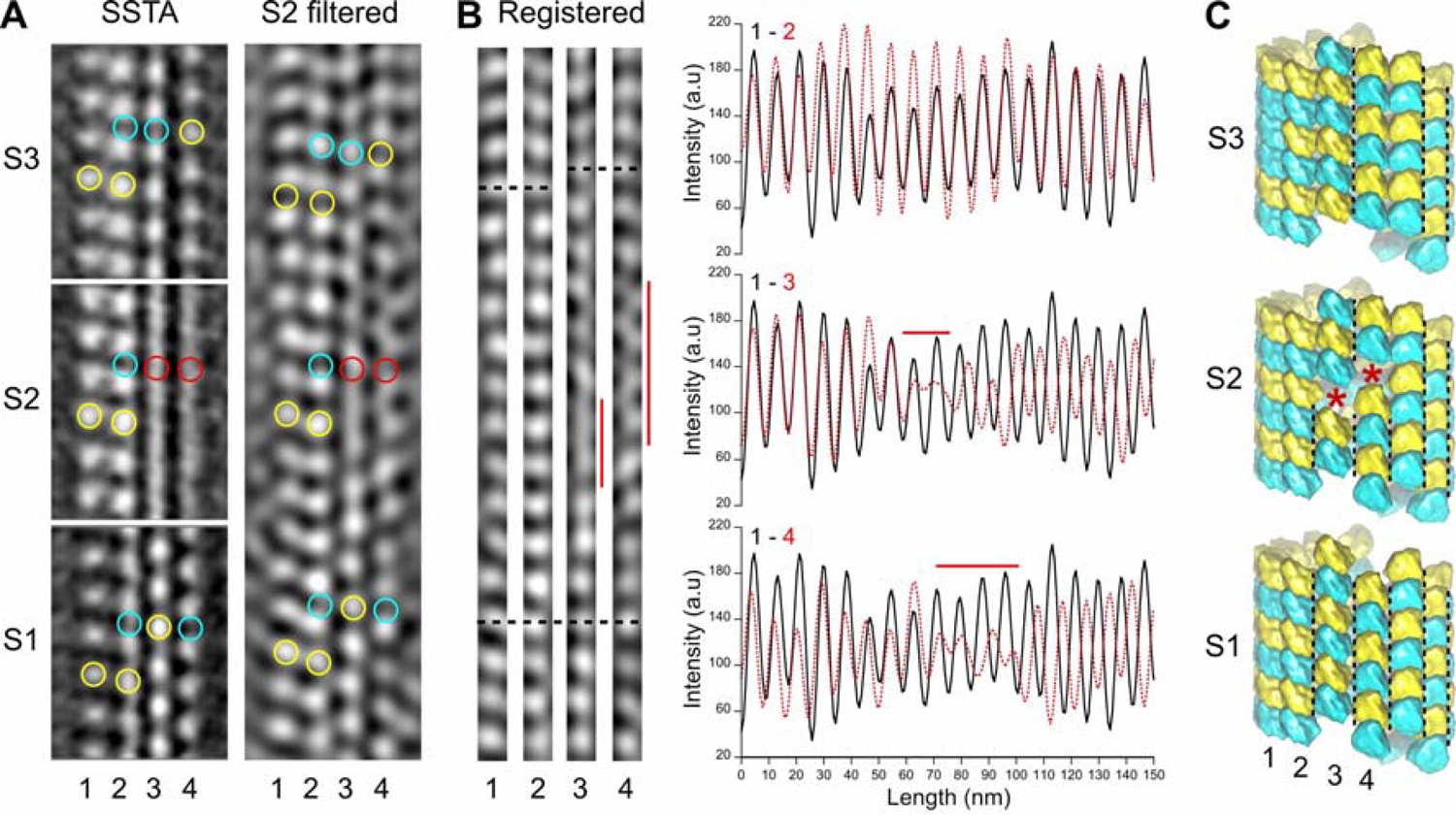
Comparison between SSTA and Fourier filtered images of transition regions. **(A)** SSTA: slices through the sub-tomogram averages of segments S1 to S3 in Figure 3B (*left*). The contrast has been inverted with respect to the original tomogram to represent protein densities as white. Yellow open circles have been placed on kinesin-motor domain densities, cyan open circles in between them, and red open circles on aberrant densities in S2. S2 filtered: slice through the filtered tomogram of the S2 region (*right*). The change in lattice seam number from S1 to S3 is clearly visualized in the S2 region. **(B)** Protofilaments 1 to 4 in (A) have been extracted from the filtered image and put into register (Registered, *left*). They remain in phase (bottom dotted line) until the densities in protofilaments 3 and 4 becomes fuzzy (vertical red lines). After theses transition zones, the kinesin-motor domain periodicity in protofilaments 3 and 4 becomes out of phase with respect to that in protofilaments 1 and 2 (upper dotted lines). These changes in kinesin-motor domain periodicity are confirmed in the line plots of the protofilaments (*right*). While the kinesin-motor domain periodicity in protofilaments 1 and 2 remain perfectly in phase (*upper graph*), it becomes out of phase for protofilaments 3 and 4 after the transition zones (*middle and bottom graphs*). **(C)** Schematic representation of the αβ-tubulin heterodimer organization in segments S1 to S3. The transition from 5 seams in S1 to 3 seams in S3 requires an offset of at least one monomer (red stars) in the protofilaments 3 and 4 of S2. Black dotted lines highlight the seams in each segment.

### Lattice type transitions involve the formation of holes within microtubules

Direct visualization of holes within microtubules self-assembled at high tubulin concentration (40 µM) in the presence of GTP was hampered by the high background generated by free tubulin in solution. In addition, the low magnification used to analyze long stretches of the microtubules was at the detriment of resolution. To improve the quality of the raw cryo-electron tomograms, we used GMPCPP to assemble microtubules at a lower tubulin concentration (10 µM), and acquired single-axis tilt series at a magnification of 50 000 X. SSTA was performed on kinesin-motor domains decorated GMPCPP-microtubules suitably oriented with respect to the tilt axis, in order to localize transition regions and to visualize corresponding holes in their lattice. The microtubule shown in Figure 5A transitioned from 1 to 3 seams as demonstrated by SSTA. Visualization of the microtubule in the raw tomogram reveals a transition from a B- to an A-lattice organization in the 3 protofilaments located at its lower surface (Figure 3B), as assessed by the diffraction patterns of the corresponding regions, and after filtration of the equatorial and 8 nm^-1^ layer lines. Enlargement of the central region (Figure 3C) shows a hole of one subunit’s size in the middle protofilament (2) that accounts for the change in lattice organization at this location. In addition, the first protofilament (1) displays a gap of one dimer’s size, although we cannot exclude that this results from an absence of kinesin-motor domain. Analysis of 31 GMPCPP-microtubules taken on 6 tomograms using the same strategy as in the presence of GTP (Figure 5 - figure supplement 4) revealed a transition frequency of 1.2 µm^-1^ (Supplementary Table 1), i.e., ∼3 fold lower than microtubules assembled in the presence of GTP. However, since we used different tubulin concentrations, i.e., 10 µM and 40 µM in the presence of GMPCPP and GTP, respectively, we cannot exclude a concentration dependent effect on the lattice type transition frequency.

**Figure 5.**
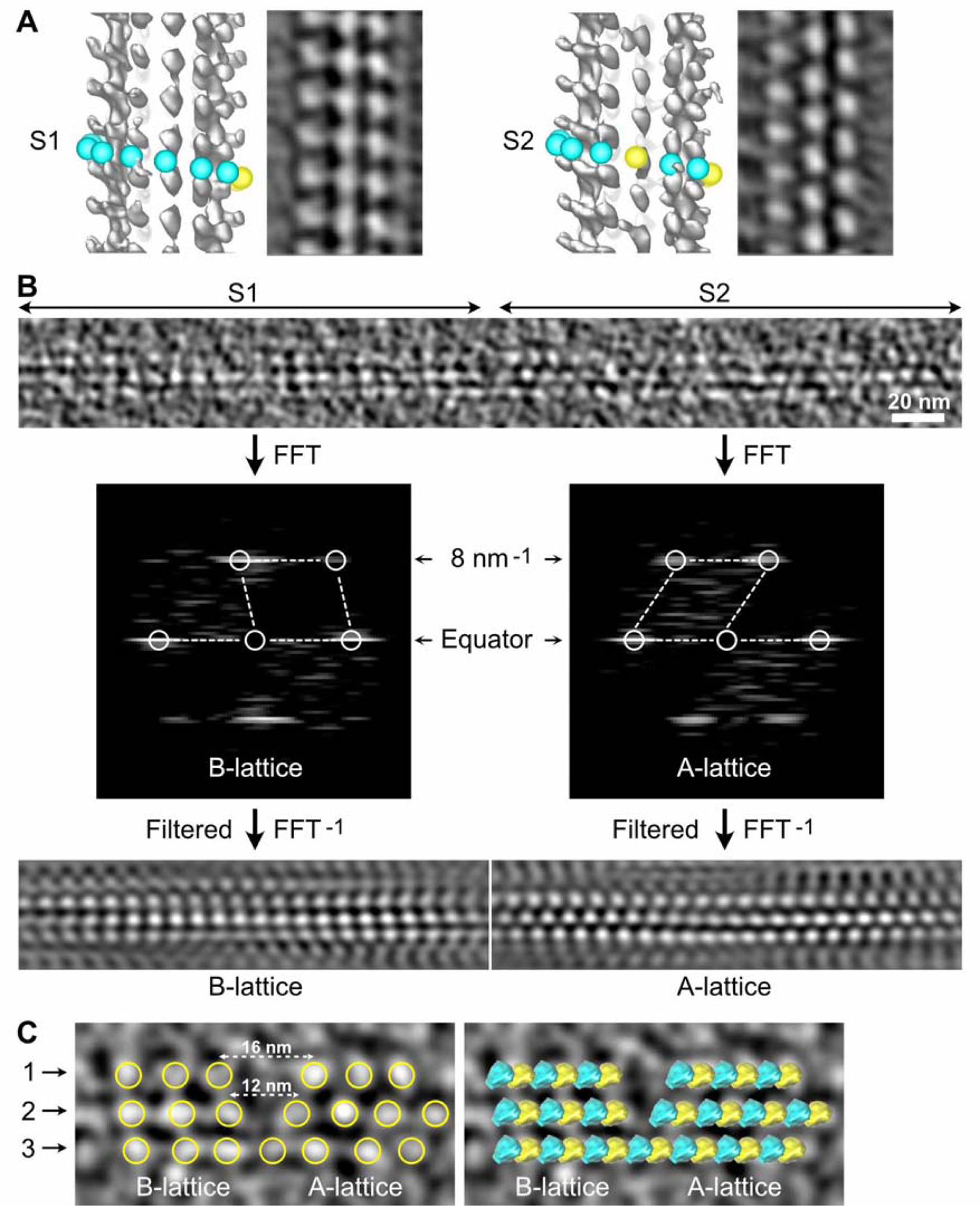
Direct visualization of holes within microtubules. **(A)** Sub-tomogram average segments before (S1) and after (S2) a lattice type transition in a GMPCPP-microtubule. For each segment, the isosurface of the full volume (*left*) and a slice through the sub-tomogram average (*right*) are displayed. The contrast has been inverted to represent protein density as white. S1 and S2 contain 1 and 3 seams, respectively. **(B)** Z-projection of 20 slices at the surface of the microtubule that encompasses S1 and S2 (*top*) with their associated Fourier transforms (*middle*) and filtered versions of the corresponding regions after selection of the equatorial and 8 nm^-1^ layer lines (*bottom*). The 3 protofilaments in S1 and S2 are organized according to a B- and an A-lattice, respectively. **(C)** Enlarged central region of the microtubule in (B). Yellow open circles have been placed on the kinesin densities (*left*), showing a gap of one subunit in protofilament 2, and possibly of a dimer in protofilament 1, although an absence of kinesin-motor domain at this location cannot be excluded. Tubulin heterodimers have been placed at the corresponding location (*right*) to highlight their change in organization at the transition region.

### Methodological artefacts limit the visualization of holes within microtubules in raw cryo-electron tomograms

During this study, we found strong limitations to the observation of holes within microtubules in raw tomograms. First, the transition regions had to be located at the top or bottom surface of the microtubule with respect to the electron beam, since edges were severely smoothed due to the lack of data at high angle that elongate densities in this direction (Figure 6A-B). This artefact is inherent to electron tomography, limiting the search of holes within microtubules in raw tomograms. A second severe artefact commonly encountered, especially in thin ice layers, was denaturation of kinesin heads at the air-water interface (Figure 6C-D). This artefact shows up as a diminution of the kinesin-motor domain density, whose periodical arrangement can only be recovered after SSTA (Figure 6D). This analysis clearly showed that SSTA remains compulsory to localize changes in lattice type organization within individual microtubules, and thus visualize the corresponding holes in regions suitably oriented with respect to the tilt axis and not in interaction with the air-water interfaces.

**Figure 6.**
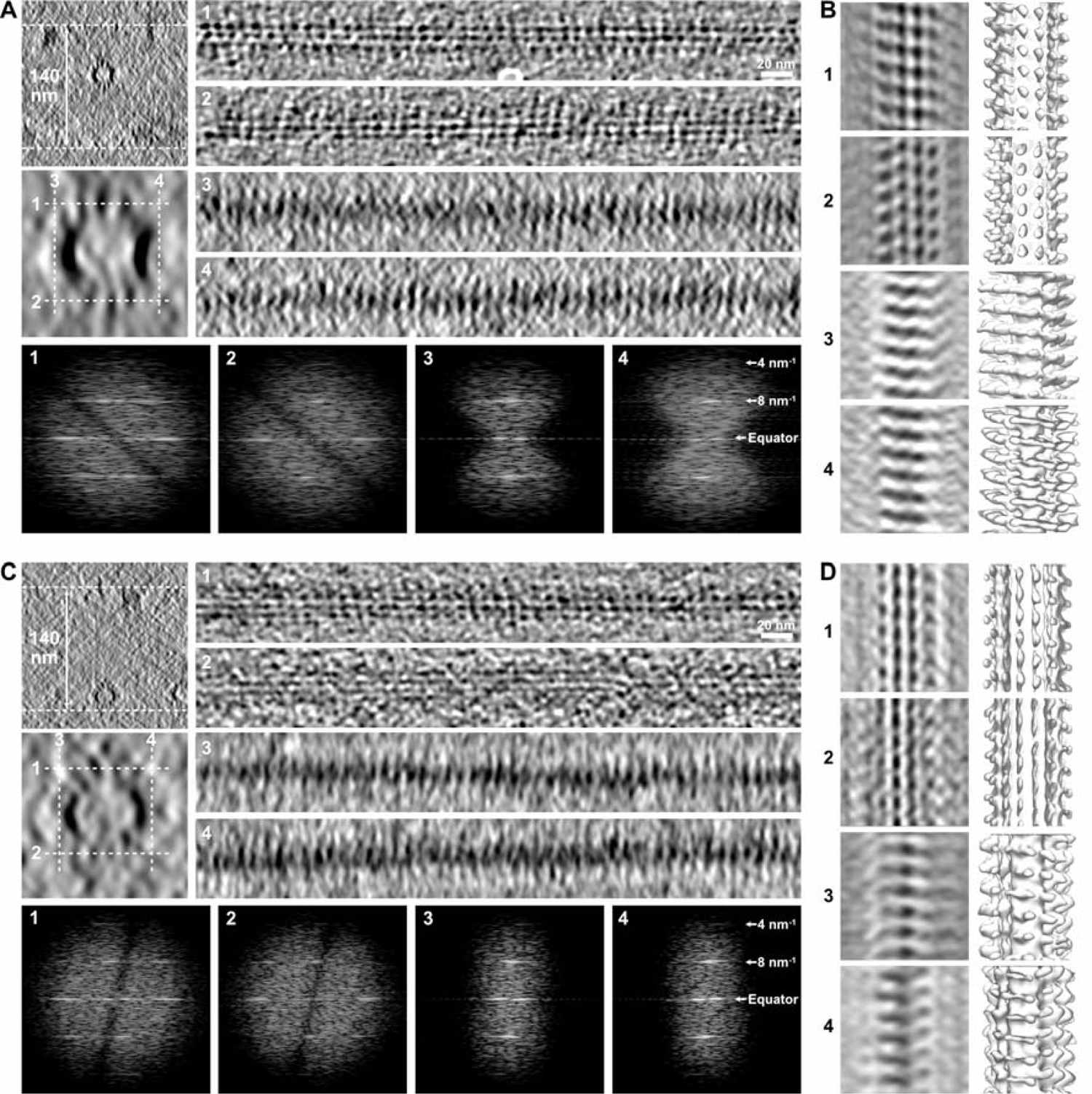
Limitations in the visualization of holes in raw tomograms. **(A)** Microtubule embedded in a ∼140 nm thick ice layer (*top left*). Longitudinal sections (averages of 20 slices, *right*) were performed at the top (1), bottom (2), left (3) and right (4) of the microtubule at positions indicated by white dotted lines in the enlarged view of the microtubule (*middle left*, average of 50 slices). Kinesin-motor domain densities can be individualized on the top (1) and bottom (2) sections, but not on the edges of the microtubules (3, 4) due to the elongation of densities in Z as a consequence of missing data at high angle. The Fourier transforms of the corresponding segments (*bottom*) shows that the 8 nm^-1^ periodicity of the kinesin-motor domains remains present in all views. **(B)** Sub-tomogram average of the microtubule in (A) over 18 kinesin-motor domain repeats. Sections (*left*) and isosurfaces (*right*) of the microtubule are displayed in correspondence to the longitudinal sections in (A). The kinesin-motor domain position is clearly observed on the top (1) and bottom (2) surfaces, and can be recovered on the microtubule edges after SSTA (3, 4). **(C)** Microtubule in the same tomogram as in (A) interacting with the air-water interface (*top left*). Kinesin-motor domain densities can be well discerned on the longitudinal sections (*right*) of the top surface facing the solution (1), but are almost indiscernible on the bottom surface that interacts with the air-water interface (2) and on the edges (3, 4). Fourier transforms (*bottom*) of the corresponding segments show that the periodicity of the kinesin-motor domains is still present, even on the damaged surface (2). **(D)** Sub-tomogram average of the microtubule in (C) over 18 kinesin-motor domain repeats. SSTA allows recovery of the kinesin-motor domain densities in all surfaces, including the one that interacts with the air-water interface (2).

### Lattice type transitions occur in a cytoplasmic environment

Next, we wondered whether the formation of holes was an intrinsic property of tubulin polymerization and if such microtubule lattice defects were also present in a cellular context. Decorating microtubules with kinesin-motor domains in cells remains challenging, since it involves removing of the cell membrane with detergents, adding kinesin-motor domains, and obtaining specimens thin enough to be analyzed by electron microscopy (Kikkawa et al., 1994; McIntosh et al., 2009). To overcome these difficulties and allow the analysis of a large data set of cytoplasmic microtubules, we took advantage of the open cellular system constituted by metaphase-arrested *Xenopus* egg cytoplasmic extracts (Gibeaux & Heald, 2019). Microtubule assembly was triggered using either DMSO (Sawin & Mitchison, 1994) or a constitutively active form of Ran (RanQ69L, (Carazo-Salas et al., 1999)) to control for possible effects of DMSO. Cryo-fluorescence microscopy was initially used to optimize the density of microtubule asters onto electron-microscope grids (Figure 7A). For structural analysis, kinesin-motor domains were added to fluorescent label-free cytoplasmic extracts right before vitrification (Figure 7 - figure supplement 1D), and specimens were imaged using dual-axis cryo-electron tomography (Figure 7B, Video 4) followed by SSTA.

**Figure 7.**
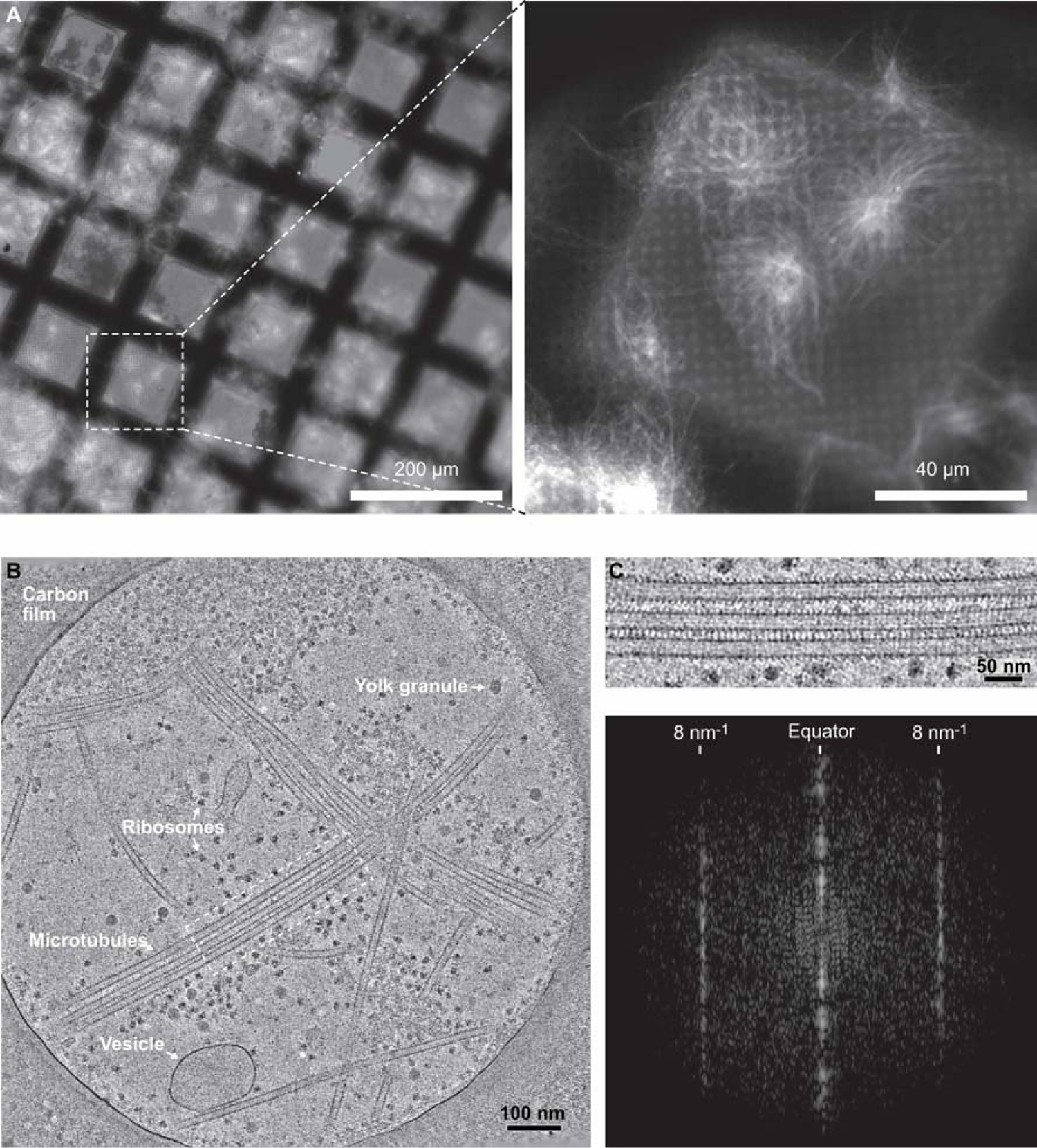
Cryo-electron tomography of microtubules decorated by kinesin-motor domains in *Xenopus* egg cytoplasmic extracts. **(A)** Cryo-fluorescence images of microtubules assembled in a cytoplasmic extract prepared from *Xenopus* eggs. Microtubules assembled in the presence of rhodamine-tubulin and plunge-frozen on an EM grid were imaged using fluorescence microscopy at liquid nitrogen temperature. Left: 10 X objective, right: 50 X objective. The white dashed square on the 10 X image indicates the field of view of the 50 X image. **(B)** Average of 30 slices in Z through a cryo-electron tomogram. The thin layer of cytoplasm spans a 2 µm diameter hole of the carbon film. The main visible features are ribosomes, vesicles, yolk granules and microtubules decorated by kinesin-motor domains. **(C)** *Top*, enlargement of the dotted rectangular region in (B) (Video 4). *Bottom*, Fourier transform of the top image showing strong layer lines at 8 nm^-1^ corresponding to the kinesin-motor domain repeat along the microtubules.

The vast majority of the microtubule segments were organized according to 13 protofilaments, three-start helices in a B-lattice configuration with one single seam (Figure 8 - figure supplement 5, Supplementary Table 2). Yet, lattice type transitions were observed in six cases over the 64 microtubules analyzed in the DMSO sample (Figure 8 - figure supplement 5, MT2, MT5, MT14, MT18, MT28, MT56). Similarly, 2 lattice type transitions were observed over 15 microtubules analyzed in the Ran sample (Figure 8 - figure supplement 6, MT4, MT10), showing that the presence of transitions was independent of the method used to trigger microtubule aster formation. The transition lattice type frequencies were ∼0.1 µm^-1^ (Supplementary Table 1), i.e., at least one order of magnitude less than with microtubules assembled from purified tubulin in the presence of GMPCPP and GTP, respectively. Strikingly, these transitions systematically involved a lateral offset of the seam by one protofilament (Figure 8, Video 5). In addition, variations in protofilament and helix-start numbers were also observed such as 12_2, 12_3, 13_4 and 14_3 microtubule-lattice regions (Figure 9). Of note, the 12_2 and 13_4 microtubules showed a local dislocation in between two protofilaments (Video 6), which is likely a response to the excessive protofilament skewing present in these microtubules (Chrétien & Fuller, 2000). The 12_2 microtubule contained two seams (Figure 9A), while the 13_4 microtubules had no seam (Figure 9C), and hence were fully helical at the tubulin dimer level (MT7 and MT8, Figure 9 - figure supplement 5A). These observations demonstrate that changes in protofilament and/or helix-start numbers, as well as multiple seams and transitions in lattice types, occur within individual microtubules assembled in a cytoplasmic context.

**Figure 8.**
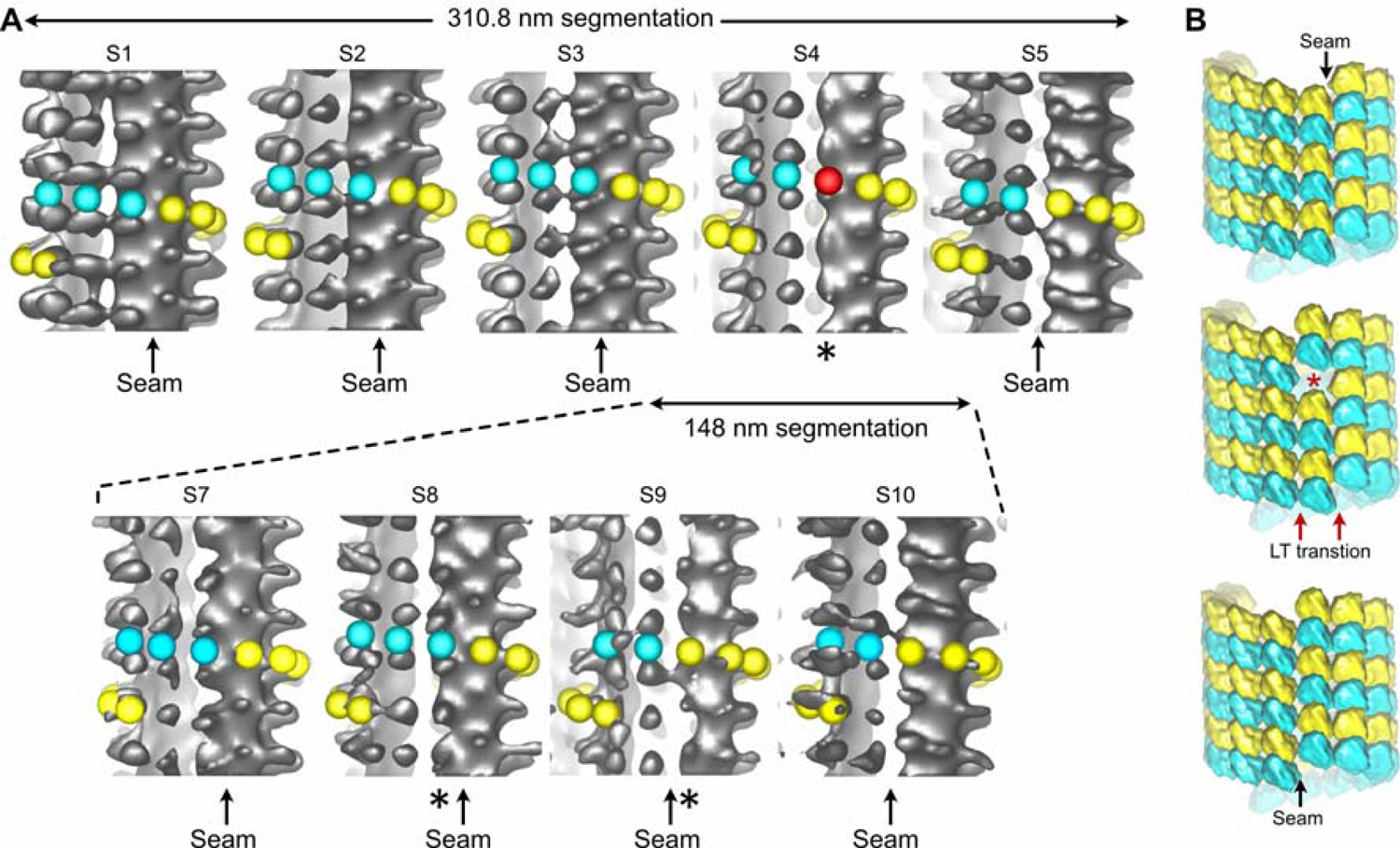
SSTA of microtubules decorated by kinesin-motor domains in *Xenopus* egg cytoplasmic extracts. **(A)** Sub-tomogram averages of five 310.8 nm long segments of a 13_3 microtubule (original length 1628 nm, *top*). S4 contains an aberrant protofilament (*), and the seam (arrow) moves laterally to the left by one protofilament from S3 to S5. The microtubule has been segmented into eleven 148 long segments (*bottom*, Figure 8 - figure supplement 5A: MT2). Only S7 to S10 are shown, corresponding to a region that encompass S3 to S5 in the 310.8 nm segmentation (Video 5). The lattice type transition occurs from S8 to S9, and no aberrant protofilament is observed in this finer segmentation. **(B)** 3D models of the tubulin lattice before (*top*), during (*middle*) and after (*bottom*) the transition. The lateral offset in seam position requires a longitudinal offset of a minimum of one tubulin subunit to account for the lattice type transition observed in (A).

**Figure 9.**
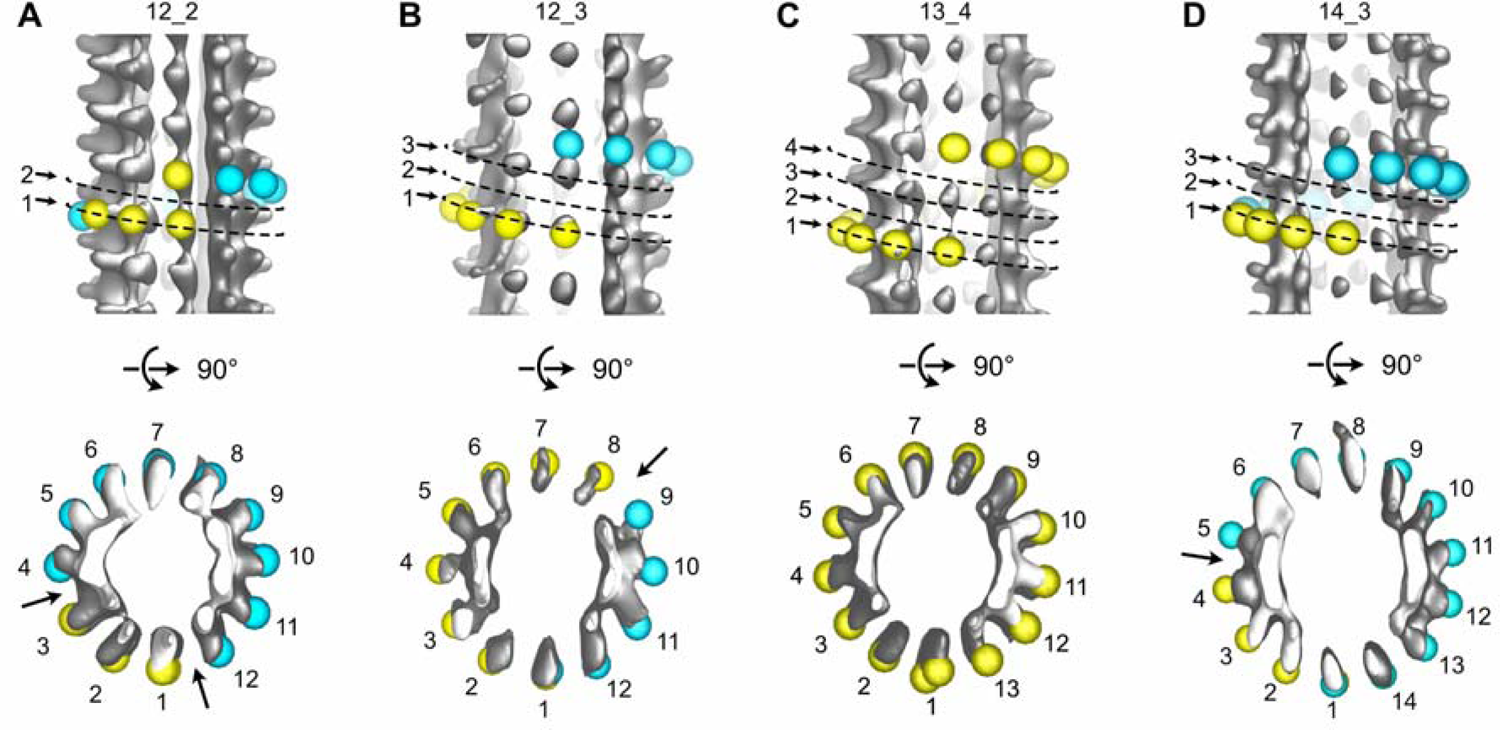
Variations in protofilament and helix-start numbers in microtubules assembled in *Xenopus* egg cytoplasmic extracts. **(A)** 12_2 microtubule with two seams (Figure 9 - figure supplement 5A: MT9). **(B)** 12_3 microtubule with a unique seam. This microtubule transitioned to a 13_3 configuration (Figure 9 - figure supplement 5D: MT62). **(C)** 13_4 microtubule with no seam. This microtubule transitioned to a 13_3 configuration (Figure 9 - figure supplement 5A: MT7). **(D)** 14_3 microtubule with one seam. This microtubule transitioned to a 13_3 configuration (Figure 9 - figure supplement 5B: MT32).

## Discussion

Here we used a segmented sub-tomogram strategy to reveal changes in lattice types within individual microtubules assembled from purified tubulin or in a cytoplasmic context, and hence holes within their lattice. Ideally, cryo-electron tomography should reveal holes in the absence of averaging. Yet, we found severe limitations that are independent of the instrument used, but that are linked to the methodology. First, missing data at high angle, whether they are taken by single- or dual-axis cryo-electron tomography, blur densities on the edges of microtubules with respect to the tilt axis (Guesdon et al., 2013). Second, we found that the interaction of the microtubules with the air-water interface diminishes the kinesin-motor domain densities, likely as a consequence of denaturation (D’Imprima et al., 2019; Klebl et al., 2022). Third, the lattice type transition frequency remains low with respect to the number of tubulin heterodimers within microtubules. It is 3.7 and 1.2 transitions every µm for microtubules assembled in the presence of GTP, and GMPCPP, respectively, and ∼1 every ∼10 µm for cytoplasmic extract microtubules. Considering that 1 µm of a 13 protofilament microtubule contains ∼1 625 dimers, this translates to one lattice type transition every 16 250 dimers, which hinders the localization of holes in raw data. Conversely, the original SSTA approach we used allows localization of lattice type transitions along individual microtubules independently of their orientation with respect to the tilt axis, and at surfaces that interact with the air-water interface. While missing data at high angle are inherent to the method of electron tomography, means to limit denaturation of proteins at the air-water interface must be found. This is critical for cryo-electron tomography, but also for single particle analysis methods where this artefact is a limiting factor to obtain high-resolution data (J. Chen et al., 2019; S. Chen et al., 2022; D’Imprima et al., 2019; Klebl et al., 2022; Li et al., 2021).

Changes in lattice types along individual microtubules could result from an imperfect annealing of shorter microtubules, a process known to occur *in vitro* (Rothwell et al., 1986). Yet, the lattice type transition frequency observed with purified tubulin would necessitate annealing of very short segments, sometime a few tens to hundreds of nm in length. The average lattice type transition frequency observed in cytoplasmic extracts could be compatible with annealing of microtubules a few µm in length. However, the fact that these transitions involved systematically a lateral seam offset of only one protofilament suggests a firm regulatory mechanism. Hence, a more plausible explanation is that these lattice discontinuities are formed during microtubule assembly (Figure 10, Video 7). At present, classical models of microtubule elongation hypothesize that tubulin engages either uniquely longitudinal interactions (Figure 10A, step 1), or both longitudinal and lateral interactions with the growing tip of microtubules (Figure 10B, step 2). A purely longitudinal elongation process (McIntosh et al., 2018) can hardly explain how microtubules can vary in terms of protofilament and/or helix start numbers as well as in lattice types, and thus how holes can arise during assembly. Conversely, to account for the presence of holes of one to a few subunits in size, it is sufficient to consider that tubulin can engage lateral interactions without longitudinal ones (Figure 10A, step 3). Gaps of an odd number of tubulin subunits will induce lattice type transitions (Figure 10A, steps 4-5), while those of an even number will induce no changes (Figure 10B). Hence, since both types of events are likely to occur, we may underestimate the presence of holes within microtubules. In addition, a finer sampling of the microtubule lattice with shorter segments could also reveal a higher hole frequency. Formation of lateral contacts without longitudinal ones at the seam region can also explain how the seam can vary in position by one protofilament (Figure 10C), since this only requires that a tubulin dimer engages homotypic lateral interactions at the seam region (Figure 10C, step 2). This event will also leave a gap of an odd number of subunits within the microtubule lattice (Figure 10C, steps 3-4).

**Figure 10.**
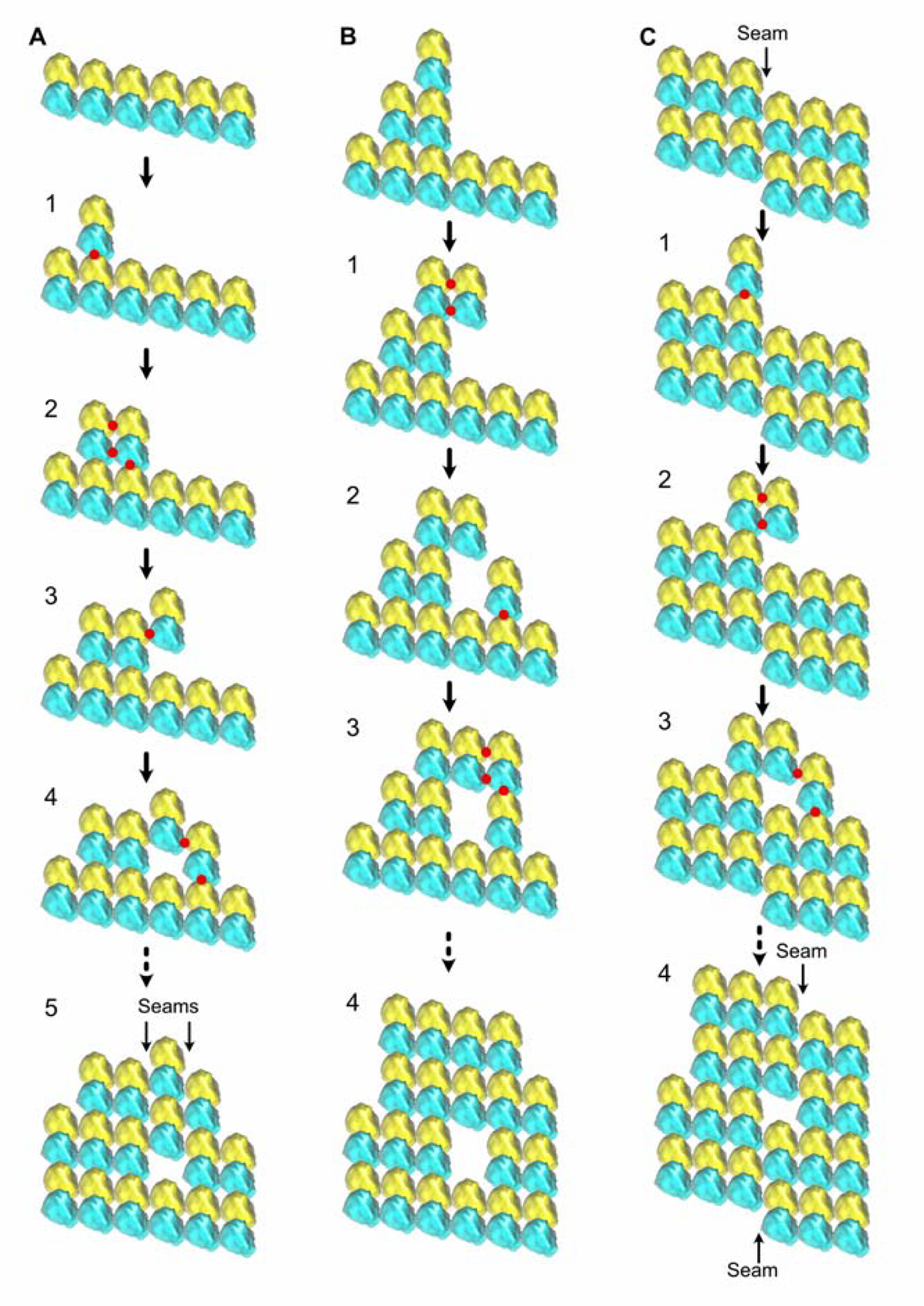
Formation of holes within microtubules during assembly. **(A)** Formation of multiple seams, red dots indicate new interactions. 1) Unique longitudinal interaction. 2) Combined lateral and longitudinal interactions. 3) Unique lateral interaction between one α-tubulin subunit of an incoming tubulin dimer and a β-tubulin subunit at the tip of the growing microtubule. 4-5) Incorporation of a hole within the microtubule lattice. Two A-lattice seams have been formed (arrows). **(B)** Incorporation of a tubulin dimer gap without change in lattice type organization. 1) Homotypic lateral interaction of an incoming tubulin dimer without longitudinal interaction. 2-5) Incorporation of a tubulin dimer gap inside the microtubule lattice. **(C)** Lateral offset of the seam by one protofilament during elongation. 1) Unique longitudinal interaction. 2) Homotypic interaction of an incoming dimer at the seam region without longitudinal contact. 3-4) Incorporation of a lattice type transition inside the microtubule wall. The seam has moved laterally by one protofilament (4), a situation systematically encountered in cytoplasmic extract microtubules.

Our current view of microtubules organized according to a perfect pseudo-helical B-lattice interrupted by a single A-lattice seam must be reconsidered. This is definitely the case for microtubules assembled from purified tubulin and has profound consequences for the interpretation of biochemical, biophysical, and structural results. For instance, 3D reconstruction studies will have to take into account the heterogeneity of the microtubule lattice to reach higher resolution (Debs et al., 2020). The lattice organization of cytoplasmic extract microtubules is more in agreement with the B-lattice, single seam model. However, exceptions are also observed such as changes in protofilament and/or helix start numbers, as well as in the location of seams within individual microtubules. Therefore, our results suggest that the formation of heterogeneous microtubule lattices is an intrinsic property of tubulin polymerization, which is firmly regulated in cells. One key regulatory factor could be the γ-tubulin ring complex (γTuRC), which imposes the 13 protofilament organization to a nascent microtubule (Böhler et al., 2021). But how this structure is preserved during microtubule elongation remains unclear, especially if one considers a two-dimensional assembly process where the lattice can vary in terms of protofilament number, helix-start number, or lattice type during elongation. Proteins of the end-binding (EB) family are other good candidates that could play a key role in regulating microtubule structure during assembly in cells. They interact with the tip of growing microtubules and bind in between protofilaments that are organized according to a B-lattice (Maurer et al., 2012); they thus may favor the formation of homotypic lateral interactions during assembly. In addition, EBs have been shown to induce the formation of 13 protofilaments, three-start helix microtubules (Manka & Moores, 2018; Vitre et al., 2008), which could also be forced to adopt a preferential B-lattice type organization. Conversely, microtubule polymerases like XMAP215, which act at growing microtubule ends (Brouhard et al., 2008), may favor lattice heterogeneities (Farmer et al., 2021). It remains to be determined whether the concerted action of different microtubule growing-end binding proteins regulate microtubule structure and dynamics in cells (Akhmanova & Steinmetz, 2008).

### Ideas and speculation

Microtubules alternate stochastically between growing and shrinking states, an unusual behavior termed dynamic instability that was discovered some 38 years ago (Mitchison & Kirschner, 1984). Although it is exquisitely regulated in cells by a myriad of microtubule associated proteins (Cleary & Hancock, 2021), it is also an intrinsic property of microtubules assembled from purified tubulin, demonstrating that it is intimately tied to tubulin assembly properties (Brouhard, 2015).

The αβ-tubulin heterodimer binds two molecules of GTP, one located between the α and β monomers at the non-exchangeable N-site, and one on the β-subunit at the longitudinal interface between heterodimers at the E-site that becomes hydrolyzed to GDP during assembly. GTP-hydrolysis destabilizes the microtubule lattice, likely weakening tubulin lateral interactions by a mechanism that remains unclear (Zhang et al., 2015). A slight delay between polymerization and GTP-hydrolysis would allow the formation of a protective GTP-cap at growing microtubule ends (Pantaloni & Carlier, 1986). The current model(s) speculate that stochastic loss of this GTP-cap induces depolymerization events known as catastrophes (Mitchison & Kirschner, 1984). However, the molecular mechanisms that lead to disappearance of the GTP-cap remain unknown. The origin of repolymerization events, termed rescues, is also unclear, but may involve tubulin molecules that have not hydrolyzed their GTP and that remain trapped inside the microtubule lattice (Dimitrov et al., 2008).

Noteworthy, the vast majority of theoretical models that have been designed so far to explain microtubule dynamic instability rely on a continuous lattice composed of B-type lattice contacts interrupted by a single seam of the A-type (Bowne-Anderson et al., 2013, 2015). Yet, exceptions to this rule have been documented over the years, essentially in microtubules assembled *in vitro* from purified tubulin. It is known that microtubules can accommodate different protofilament and helix-start numbers (Chaaban & Brouhard, 2017; Chrétien & Wade, 1991). These numbers can vary within individual microtubules (Chrétien et al., 1992), necessarily leaving holes inside their lattice (Schaedel et al., 2019; Théry & Blanchoin, 2021). Microtubules can also adopt configurations with a high protofilament skew that must be compensated by a relaxation step whose detailed mechanism remains to be described (Chrétien & Fuller, 2000). Microtubules with different numbers of seams have been described (Debs et al., 2020; des Georges et al., 2008; Howes et al., 2017; Kikkawa et al., 1994; Sosa et al., 1997), although it was not considered that both the seam number and location could vary within individual microtubules. Therefore, these previous studies and the present one indicate that the microtubule lattice is highly labile, with the ability to form different kinds of structural defects (Hunyadi et al., 2005; Rai et al., 2021).

The formation of lattice defects during microtubule polymerization must impose energetical penalties at the growing microtubule end, potentially destabilizing the protective GTP-cap if present, and hence be at the origin of catastrophes. Likewise, holes must let patches of unhydrolyzed GTP-tubulin molecules within microtubules, potentially at the origin of rescues. Hence, we propose that microtubule dynamic instability is not only driven by the nucleotide state of tubulin, but also by the intrinsic structural instability of the microtubule lattice. MAPs such as EBs and XMAP215 may exploit this structural instability to finely tune microtubule dynamics in cells.

## Materials and Methods

### Protein purification

Tubulin was isolated from porcine brain by two cycles of assembly disassembly (Castoldi & Popov, 2003), followed by a final cycle in the absence of free GTP (Ashford & Hyman, 2006). Tubulin was obtained in BRB80 (80 mM K-Pipes, 1 mM EGTA, 1 mM MgCl_2_, pH 6.8 with KOH) and stored at −80 °C before use.

The cDNA fragment encoding for the human Kif5B motor domain (residues 1 to 349) was cloned into the pET based bacterial vector PSTCm1 (Olieric et al., 2010). The protein was expressed in Rosetta2 *E. coli* cells. Cells were grown at 37 °C in LB media supplemented with 50 μg/ml kanamycin and 30 μg/ml chloramphenicol to an OD_600_ of 0.4 to 0.6. Temperature was reduced to 20 °C, the protein production was induced 20 min later with 0.5 mM IPTG (isopropy-1-thio-β-galactopyranoside), and incubation was continued overnight under agitation. The cells were harvested by centrifugation for 15 min at 4 000 *g* and the cell pellets were resuspended in lysis buffer (50 mM HEPES, pH 8.0, supplemented with 10 mM imidazole, 10 % glycerol, 0.1 mM ADP, 2 mM beta-mercaptoethanol, and one cOmplete EDTA free proteases inhibitor cocktail tablet per 50 mL buffer). The cells were lysed on ice per ultrasonication and lysate clearing was performed by centrifugation, 30 min at 24 000 *g*. The resultant supernatant was filtered using a 0.45 µm filter and the protein was subsequently purified by IMAC on a 5 ml HP HisTrap column (GE Healthcare) according to manufacturer’s information. The eluted protein from this affinity step was concentrated and further purified by gel filtration on a HiLoad® 16/600 Superdex® 200 pg column (GE Healthcare) equilibrated in 20 mM TrisHCl, pH 7.5, supplemented with 150 mM NaCl, 0.1 mM ADP, and 2 mM DTT. The homogeneity of the recombinant Kif5B motor domain was assessed by SDS-PAGE. Fractions were concentrated, aliquoted, flash frozen into liquid nitrogen, and stored at −80 °C.

### Animals

All animal experimentation in this study was performed according to our animal use protocol APAFiS #26858-2020072110205978 approved by the Animal Use Ethic Committee (#7, Rennes, France) and the French Ministry of Higher Education, Research and Innovation. Mature *Xenopus laevis* female frogs were obtained from the CRB Xénope (Rennes, France) and ovulated with no harm to the animals with at least a 6-month rest interval between ovulations.

### *Xenopus* egg cytoplasmic extracts

Cytostatic factor-arrested (CSF) egg extracts were prepared from freshly laid eggs of *Xenopus laevis* as previously described (Good & Heald, 2018; Murray, 1991). Briefly, eggs arrested in metaphase of meiosis II were collected, dejellied and fractionated by centrifugation. The cytoplasmic layer was isolated, supplemented with 10 mg/ml each of the protease inhibitors leupeptin, pepstatin and chymostatin (LPC), 20 mM cytochalasin B, and a creatine phosphate and ATP energy regeneration mix. Vitrification of the samples for cryo-electron microscopy was performed the same day as the egg extract preparation.

### Cryo-fluorescence microscopy

To determine the optimal density of microtubule structures assembled from *Xenopus* egg cytoplasmic extracts cryo-fixed on electron microscopy grids suitable for cryo-electron tomography acquisitions, we used cryo-fluorescence microscopy. Egg extracts were supplemented with 40 ng/µg rhodamine-tubulin before microtubule assembly was conducted by addition of 5% DMSO and incubation at 23 °C, for 30 to 45 min. Reactions were then extemporaneously diluted 1:10, 1:50 or 1:100 in 1X BRB80 buffer for vitrification on an electron microscopy grid. Frozen grids were imaged within a Linkam CMS196M cryo-correlative microscopy stage mounted on an Olympus BX51 microscope equipped with a Lumencor SOLA SE U-nIR light source, UPLFLN10X/0.30 and LMPLFLN50X/0.50 objectives, and a Photometrics Prime-BSI sCMOS Back Illuminated camera. Images were acquired using the µManager acquisition software v1.4 (Edelstein et al., 2014).

### Cryo-electron tomography

Microtubules were assembled from purified porcine brain tubulin at 40 µM in BRB80, 1 mM GTP, or at 10 µM in BRB80, 0.1 mM GMPCPP, for about 1 h. Kif5B was diluted at a final concentration of 2.5 mg/ml in BRB80, 0.1 mM ATP, 1 mM GTP and 60 nM mix-matrix capped gold nanoparticles (Duchesne et al., 2008; Guesdon et al., 2016) and pre-warmed at 35°C. First, 3 µl of the microtubule sample was deposited at the surface of a glow-discharged holey carbon grid (Quantifoil R2/2, Cu200) in the temperature (35 °C) and humidity-controlled atmosphere (∼95 %) of an automatic plunge-freezer (EM-GP, Leica). Then, 3 µl of the pre-warmed kinesin motor domain suspension was added to the grid onto the sample, incubated for 30 s, and blotted manually. An additional 3 µl of the pre-warmed kinesin motor domain suspension was added to the grid, blotted with the EMGP for 2 s using Whatman grade 1 filter paper and plunged into liquid ethane.

Microtubule aster assembly was induced in *Xenopus* egg cytoplasmic extracts by adding 5% DMSO or 15 µM of the GTPase deficient Ran mutant RanQ69L purified as previously described (Helmke & Heald, 2014), and incubating at 23 °C for 30 to 45 min. Kif5B was diluted at a final concentration of 2.5 mg/ml in BRB80, 0.1 mM ATP, 1 mM GTP and 60 nM mix-matrix capped gold nanoparticles (Duchesne et al., 2008; Guesdon et al., 2016), and pre-warmed at 23 °C. A 3 µl volume of the Kif5B suspension was first deposited at the surface of a glow-discharged holey carbon grid (Quantifoil R2/2, Cu200) in the temperature (23 °C) and humidity-controlled atmosphere (∼95 %) of the EM-GP, on the side of the grid to be blotted. Right away, one volume of the extract sample was diluted 50 X in the pre-warmed Kif5B suspension, and 3 µl of this mix was deposited on the other side of the grid. The grid was blotted from the opposite side of the sample with the EM-GP for 4 s using Whatman grade filter 4 and plunged into liquid ethane.

For dual-axis cryo-electron tomography, specimen grids were transferred to a rotating cryo-holder (model CT3500TR, Gatan), and observed using a 200 kV electron microscope equipped with a LaB_6_ cathode (Tecnai G^2^ T20 Sphera, FEI). Images of microtubules assembled from purified tubulin were recorded on a 4k x 4K CCD camera (USC4000, Gatan) in binning mode 2 and at a nominal magnification of 29 000 X, providing a final pixel size of 0.79 nm. Images of microtubules assembled in *Xenopus* egg extracts were recorded on a 4K x 4k CMOS camera (XF416, TVIPS) in binning mode 2 and at a nominal magnification of 25 000 X and 29 000 X, providing final pixel sizes of 0.87 nm and 0.74 nm, respectively. Pixel sizes were calibrated using TMV as a standard (Guesdon et al., 2016). Dual-axis cryo-electron tomography data were acquired as previously described (Guesdon et al., 2013). Briefly, a first tilt series of ∼40 images was taken in an angular range of ∼±60° starting from 0° and using a Saxton scheme. The specimen was turned by a ∼90° in plane rotation at low magnification and a second tilt series was taken on the same area using parameters identical to the first series. Tomograms were reconstructed using the Etomo graphical user interface of the IMOD program (Kremer et al., 1996; Mastronarde, 1997). Tilt series were typically filtered after alignment using a low pass filter at 0.15 cycles/pixels and a sigma of 0.05. Tomograms were reconstructed in 3D using the SIRT-like filter of Etomo with 15 equivalent iterations. Dual-axis cryo-electron tomograms were converted to bytes before further processing.

For single-axis cryo-electron tomography, specimen grids were transferred to a dual grid cryo-transfer holder model 205 (Simple Origin). Data were acquired on a 4K x 4k CMOS camera (XF416, TVIPS) in binning mode 1 and at a nominal magnification of 50 000 X, providing a final pixel size of 0.21 nm. Typically, 40 images were taken in an angular range of ∼±60° starting from 0°, or using a symmetric electron dose scheme (Hagen et al., 2017). To localize holes within microtubules by SSTA, tomograms were subsequently binned by 4 to provide a final pixel size of 0.83 nm.

### Sub-tomogram averaging

Sub-tomogram averages were calculated using the procedure described online (https://bio3d.colorado.edu/RML_2017/2017_IMOD_PEET_Workshop/Lab_Instructions/L8 Microtubules.pdf). Briefly, a first model was created by following individual protofilaments in cross section using the slicer tool in IMOD. Usually, ∼50 electronic slices were averaged to reinforce the contrast. A second model was next extrapolated from the first one to mark the microtubule center at the same point positions. Then, a third model was calculated from the previous ones with points spaced every ∼8 nm, and a motive list containing Euler angles of each sub-volume with respect to the chosen reference was created. Sub-volumes of ∼40 pixels^3^ were extracted at each point position using the graphical user interface of the PEET program (Nicastro et al., 2006). Registration of the microtubule segments was performed by cross-correlation, limiting rotational angular searches around the microtubule axis to about half the angular separation between protofilaments. Other angles were set to take into account variations of microtubule curvature in the X, Y and Z directions. Segmented sub-tomogram averaging was performed using a new routine (splitIntoNsegments) implemented into the PEET program version 1.14.1. This routine splits the initial model and motive list into *N* segments of equal size and creates sub-directories for each segment. Sub-tomogram averages are calculated for each segment, using the original sub-tomogram average parameters of the whole microtubule as a template.

### Image analysis and model building

Sub-tomogram averages were inspected using the isosurface panel of IMOD. Four scattered models were created. Model 1 was used to mark the kinesin motor domain densities (yellow spheres), model 2 to mark the absence of densities (cyan spheres), model 3 aberrant densities (red spheres), and model 4 the microtubule center. Spheres from model 1 to 3 were placed along the *S*-start lateral helices. The 4^th^ last model was enlarged to cross the kinesin motor domain densities in order to place the other spheres at a same radius. The number of protofilaments and the different lateral contacts (A, B and undefined lateral contacts) were retrieved from these models.

## Acknowledgments

Cryo-electron microscopy data were acquired on the Microscopy Rennes imaging center platform (Biosit, Rennes, France), member of the national infrastructure France-BioImaging (FBI) supported by the French National Research Agency (ANR-10-INBS-04). *Xenopus laevis* eggs were obtained from the Centre de Ressources Biologique Xénopes, Université de Rennes 1, Rennes, France. Porcine brains were kindly provided by Y. Drillet, Cooperl Arc Altantique, Lamballe France. Tobacco Mosaic Virus was kindly provided by T. Candresse, UMR 13332 Biologie du Fruit et Pathologie, INRAE and University of Bordeaux, Villenave d’Ornon, France. Video 7 was designed by A. Kawska, Illuscienta, Paris, France. This work was supported by two French National Research Agency grants (ANR-16-CE11-0017-01 to D.C. and M.O.S., and ANR-18-CE13-0001-01 to D.C.), a Swiss National Science Foundation grant (310030_192566 to M.O.S.) and a Human Frontier Science Program grant (CDA00019/2019-C to R.G.).

---

**Figure supplement 1.**
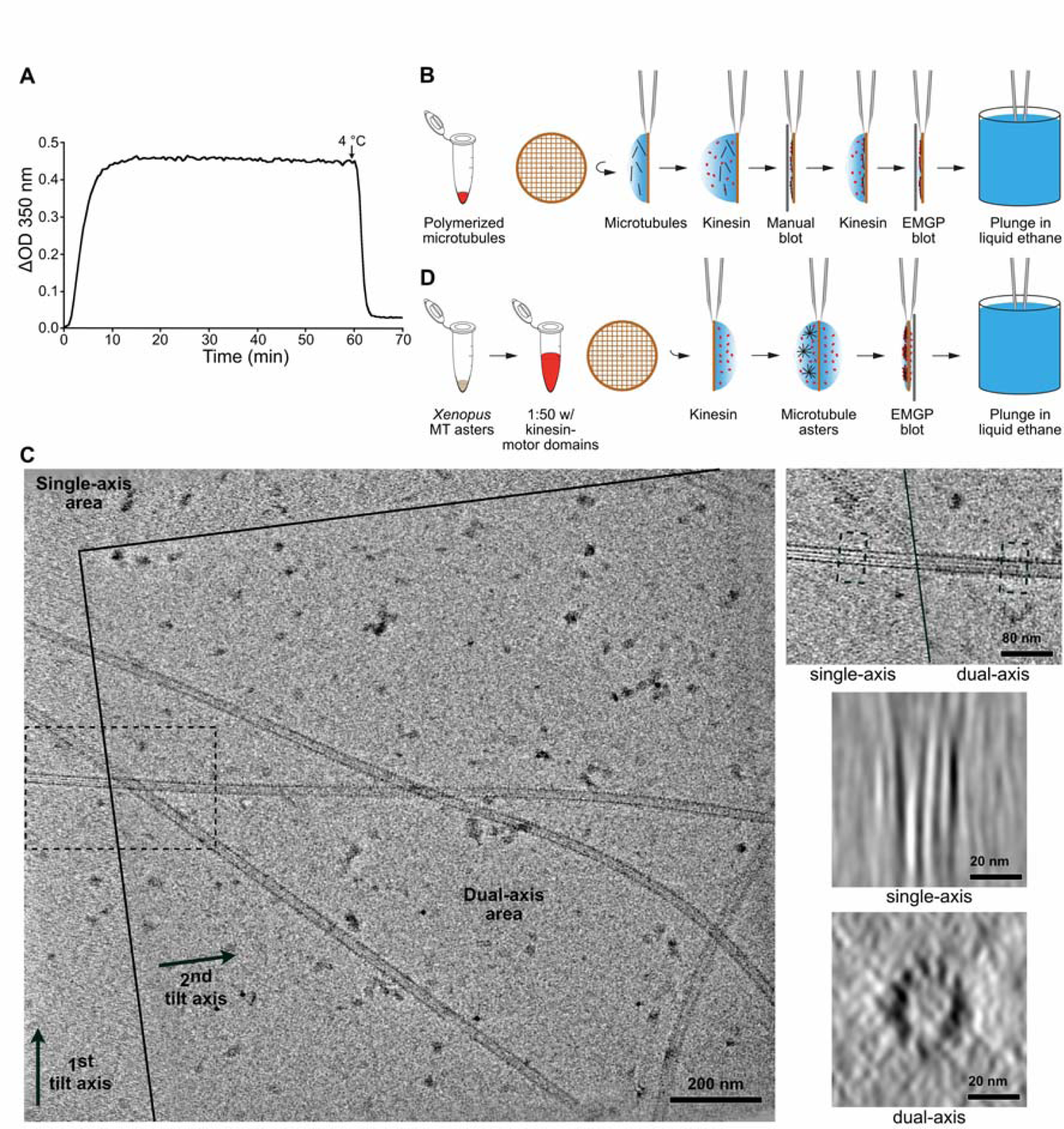
Preparation of microtubules for cryo-electron tomography. **(A)** Microtubule assembly at 40 µM tubulin in BRB80, 1 mM GTP, 35 °C. Variation of optical density (ΔOD) at 350 nm versus time (min). A cooling step was typically performed at the end of the experiment to assess the presence of aggregates evidenced by the value of the baseline at 4 °C. Samples for cryo-electron microscopy were prepared after ∼1 hour of polymerization. **(B)** Decoration of microtubules assembled *in vitro* from purified tubulin with kinesin-motor domains. Kinesin decoration is performed right before vitrification of the sample into liquid ethane. **(C)** Dual-axis cryo-electron tomography. *Left*: The dual-axis area is delimited by plain lines; arrows indicate the two tilt axes. *Right top*: Enlargement of the dotted region in the overall view that encompass the single and dual axes regions of the tomogram. A microtubule oriented close to the perpendicular of the 1^st^ tilt axis spans the two regions (Video 1). *Right middle*: Average of 50 slices along the microtubule in the single-axis area. The microtubule is severely elongated in the Z direction. *Right bottom*: The same microtubule visualized in the dual-axis area. Individual protofilaments are clearly individualized. **(D)** Decoration of microtubules assembled in *Xenopus* egg cytoplasmic extracts. Microtubule asters are deposited on one side of the grid, which is subsequently blotted from the opposite side.

**Figure supplement 2.**
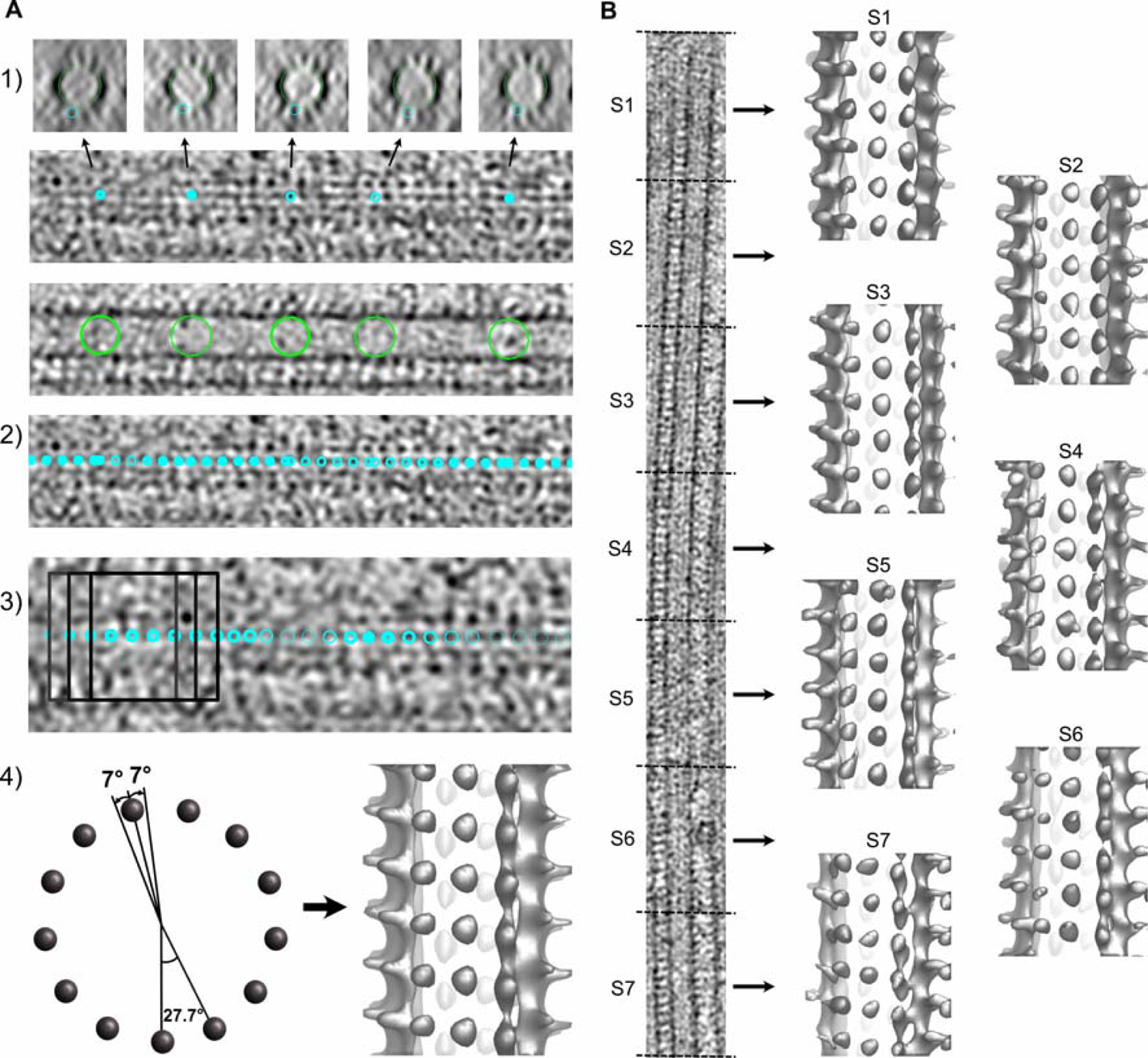
Segmented sub-tomogram averaging of microtubules decorated with kinesin motor domains. **(A)** Sub-tomogram averaging (STA). (1) A first model is created by placing model points onto individual protofilaments (small cyan circles) and at the microtubule center (large green circles). (2) A second model is extrapolated from the previous one with points spaced every ∼8 nm along the microtubule. (3) Sub-volumes (dark squares) are extracted at every point position and aligned by cross-correlation using PEET. (4) The rotational search around the microtubule axis is restricted to about half the angular separation between protofilaments (*left*). The resulting average is displayed in isosurface rendering in IMOD (*right*). **(B)** Segmented sub-tomogram averaging (SSTA). The model and motive list used to calculate the whole tomogram of the microtubule is split into shorter segments of equal dimensions and sub-tomograms are calculated for each segment (S1 to S7). The segment size is limited by the signal-to-noise ratio present in the tomograms (typically ∼160 nm with our current data).

**Figure supplement 3.**
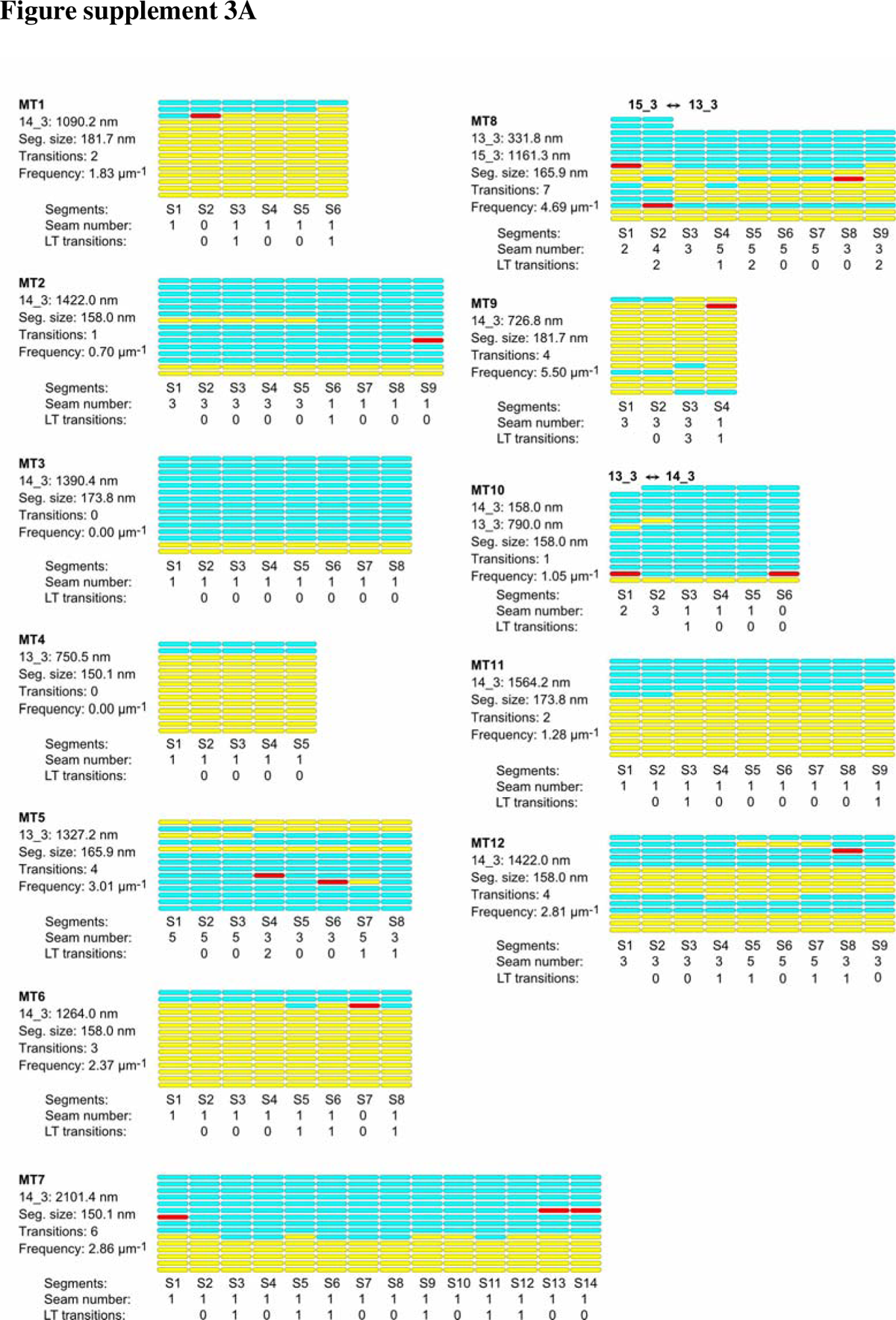
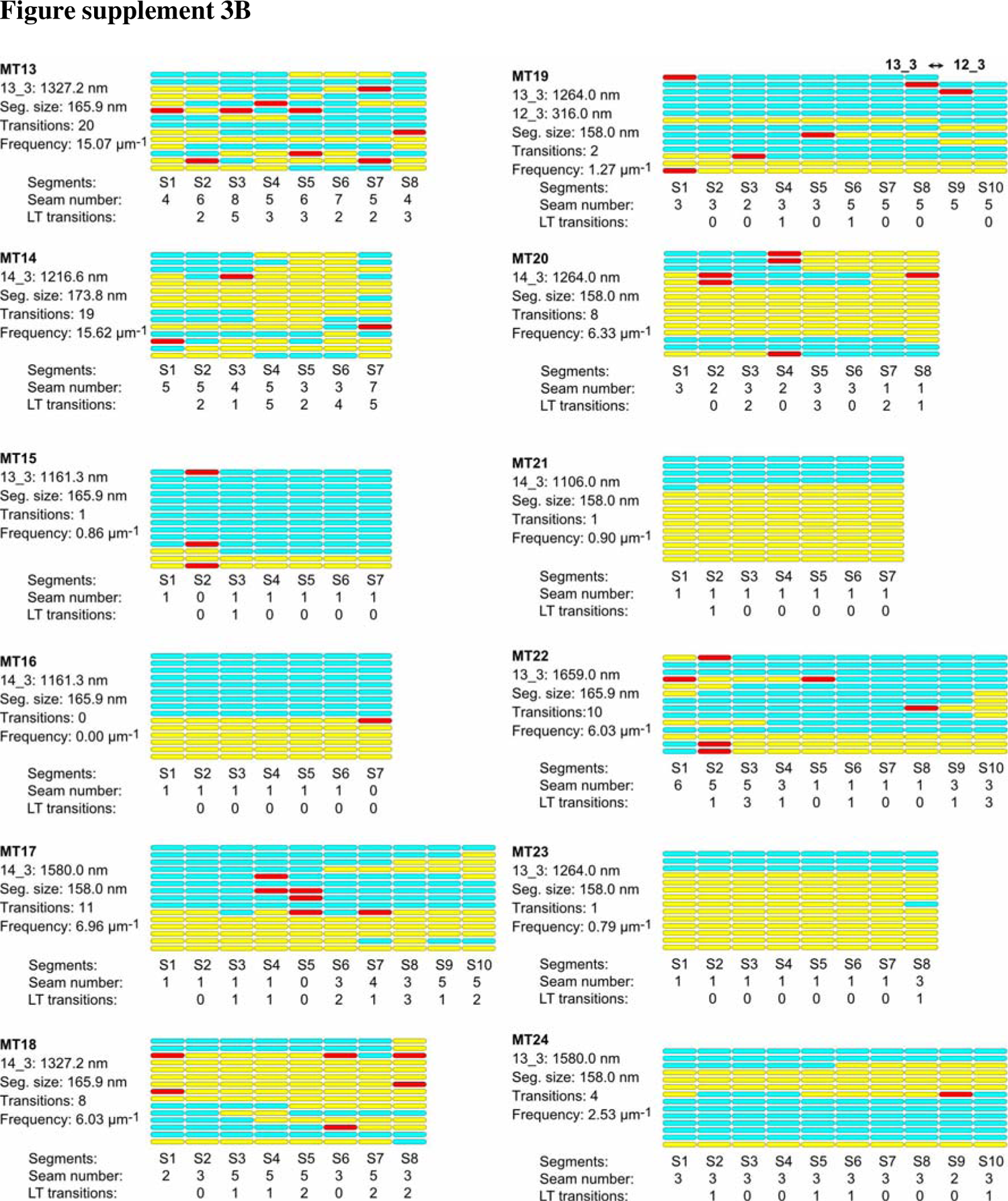
Lattice organization of microtubules assembled *in vitro* from purified porcine brain tubulin in the presence of GTP 24 microtubules (31.7 µm in total length) were analyzed on 4 tomograms acquired on 2 independent samples. Number of microtubule segments: 195; number of lateral interactions: 2664 (A type: 461, B type: 2091, ND: 112); lattice type transitions: 119; protofilament/helical-start (*N/S*) number transitions: 3.

**Figure supplement 4.**
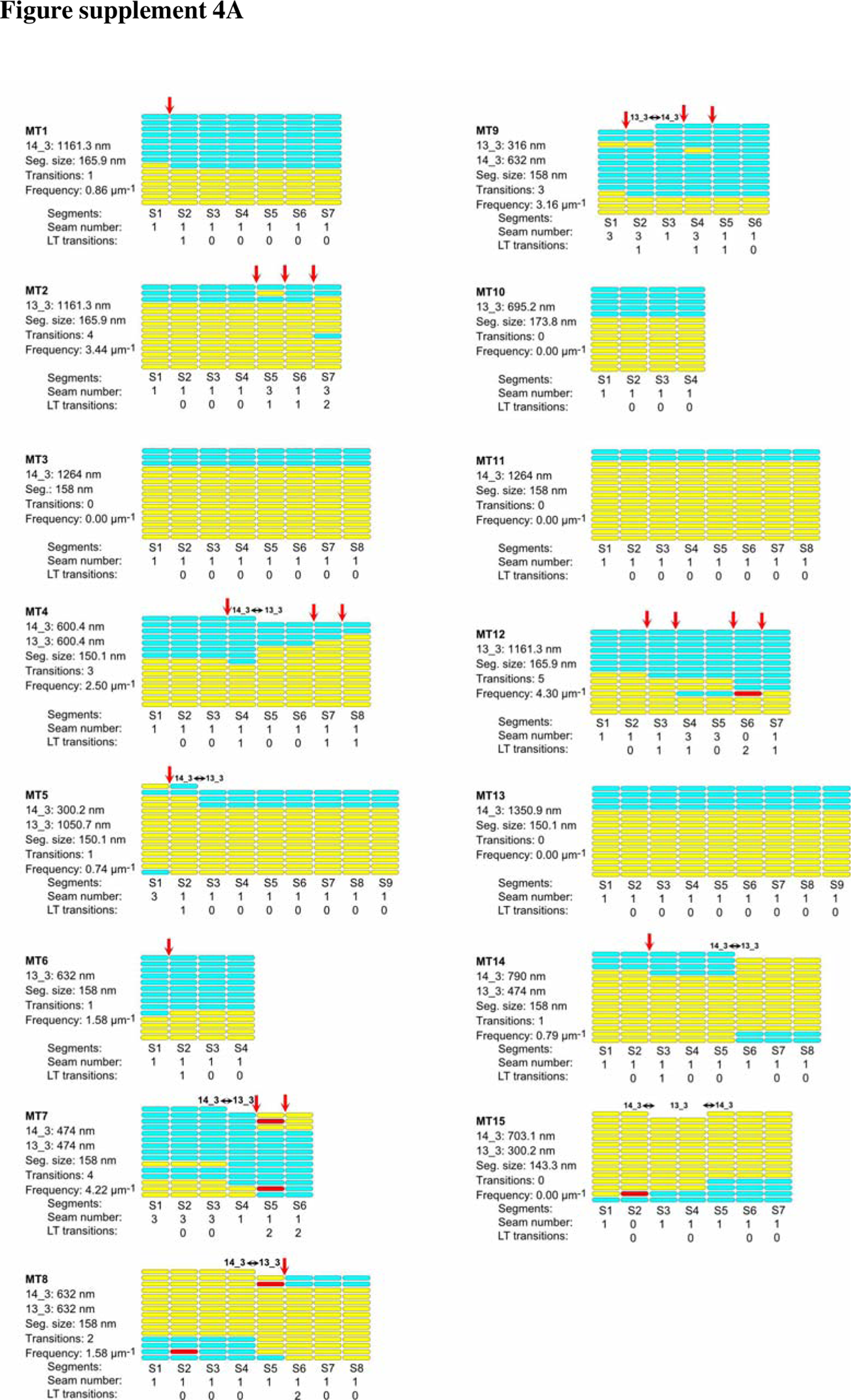
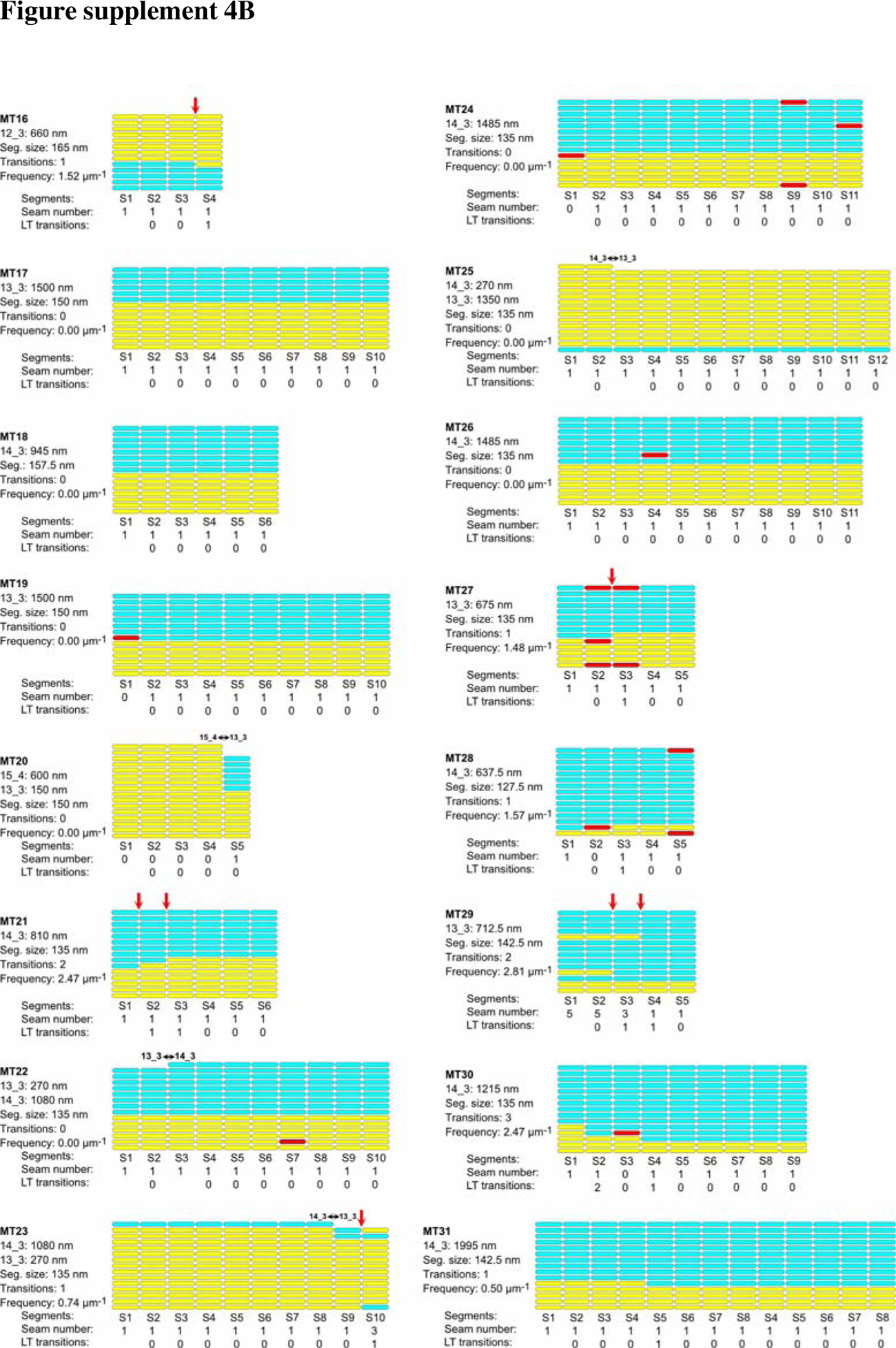
Lattice organization of microtubules assembled *in vitro* from purified porcine brain tubulin in the presence of GMPCPP 31 microtubules (35.4 µm in total length) were analyzed on 6 tomograms acquired on 2 independent samples. Number of microtubule segments: 239; number of lateral interactions: 3250 (A type: 261, B type: 2951, ND: 38); lattice type (LT) transitions: 37; protofilament/helical-start (*N/S*) number transitions: 12.

**Figure supplement 5.**
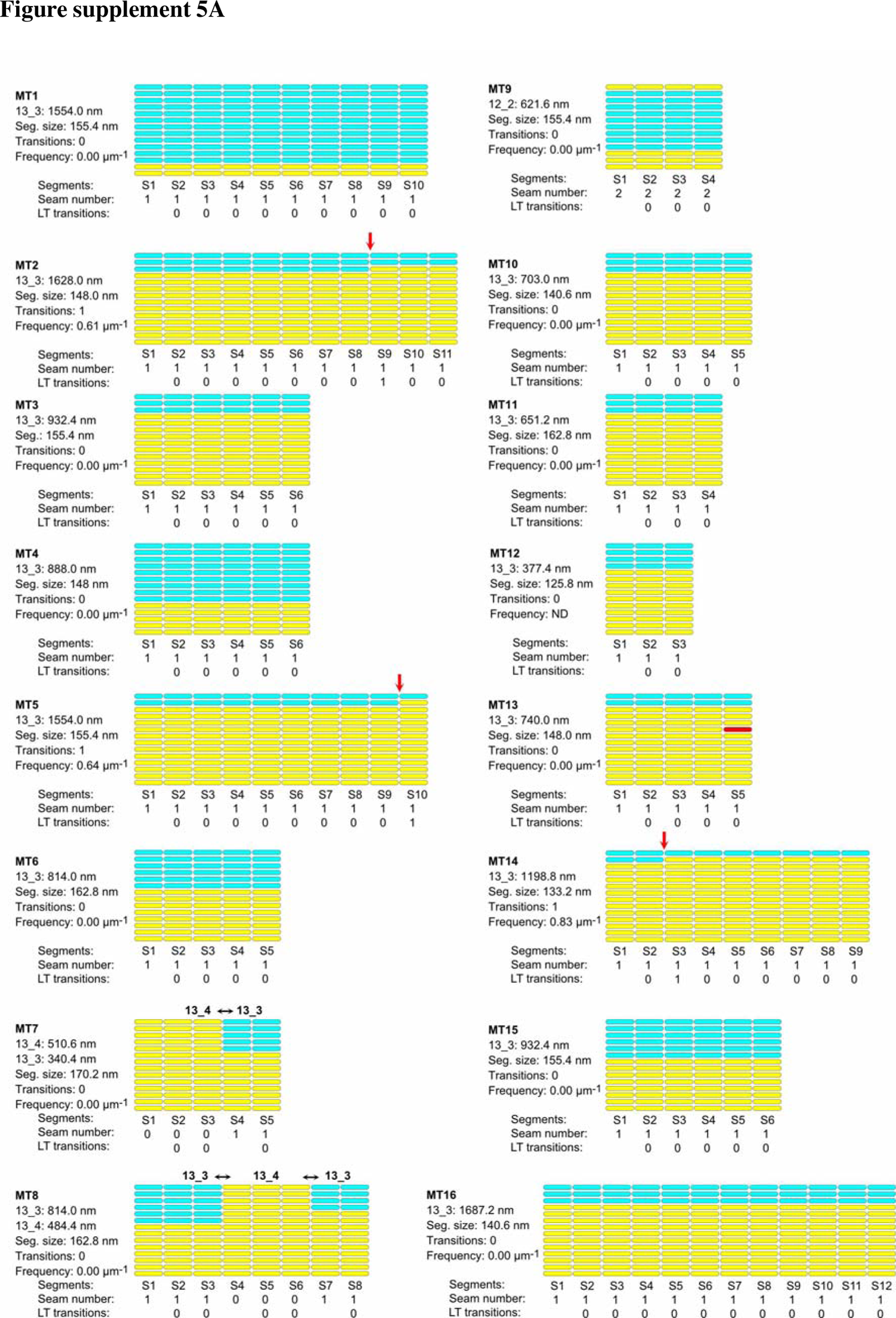
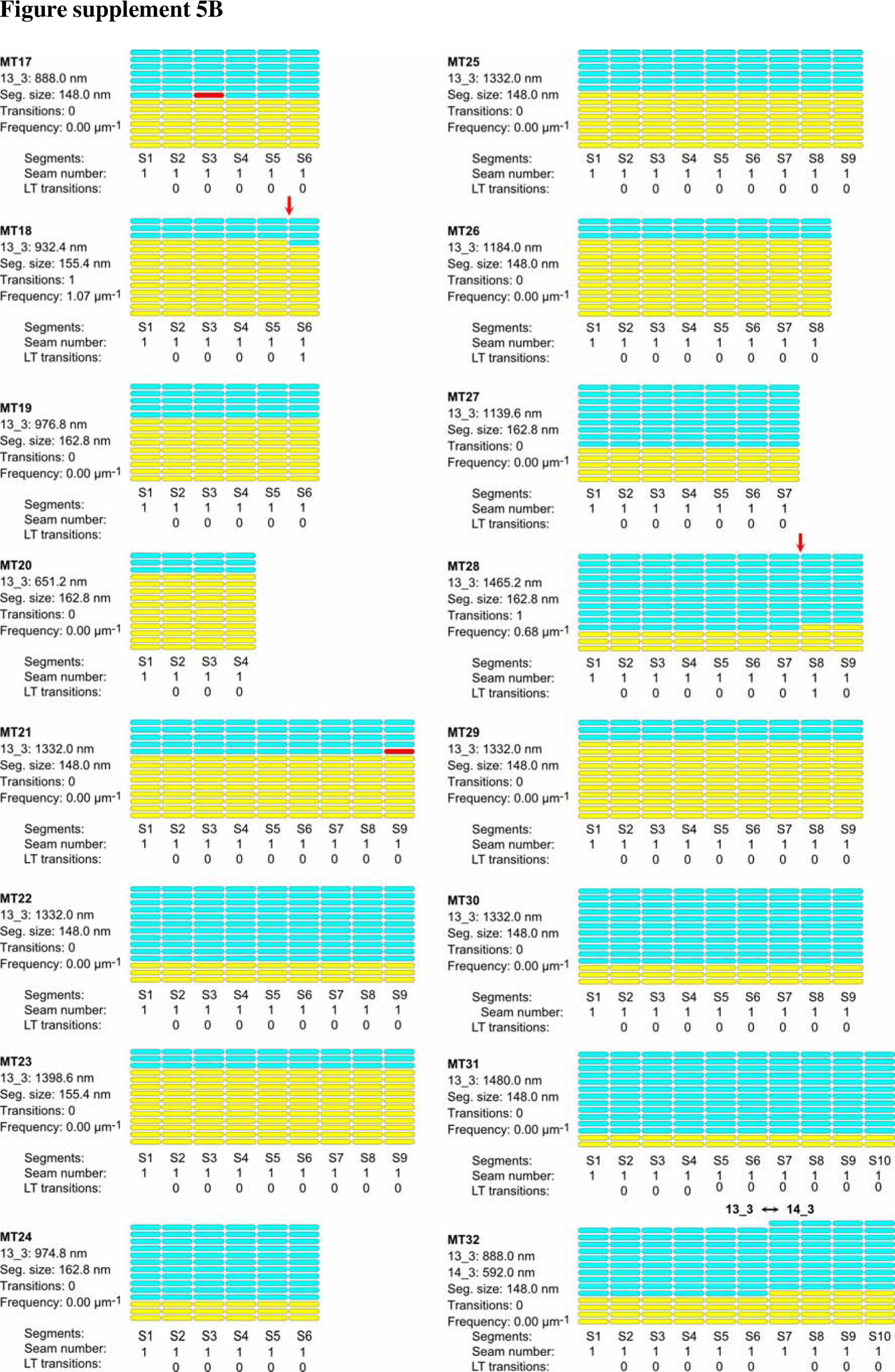
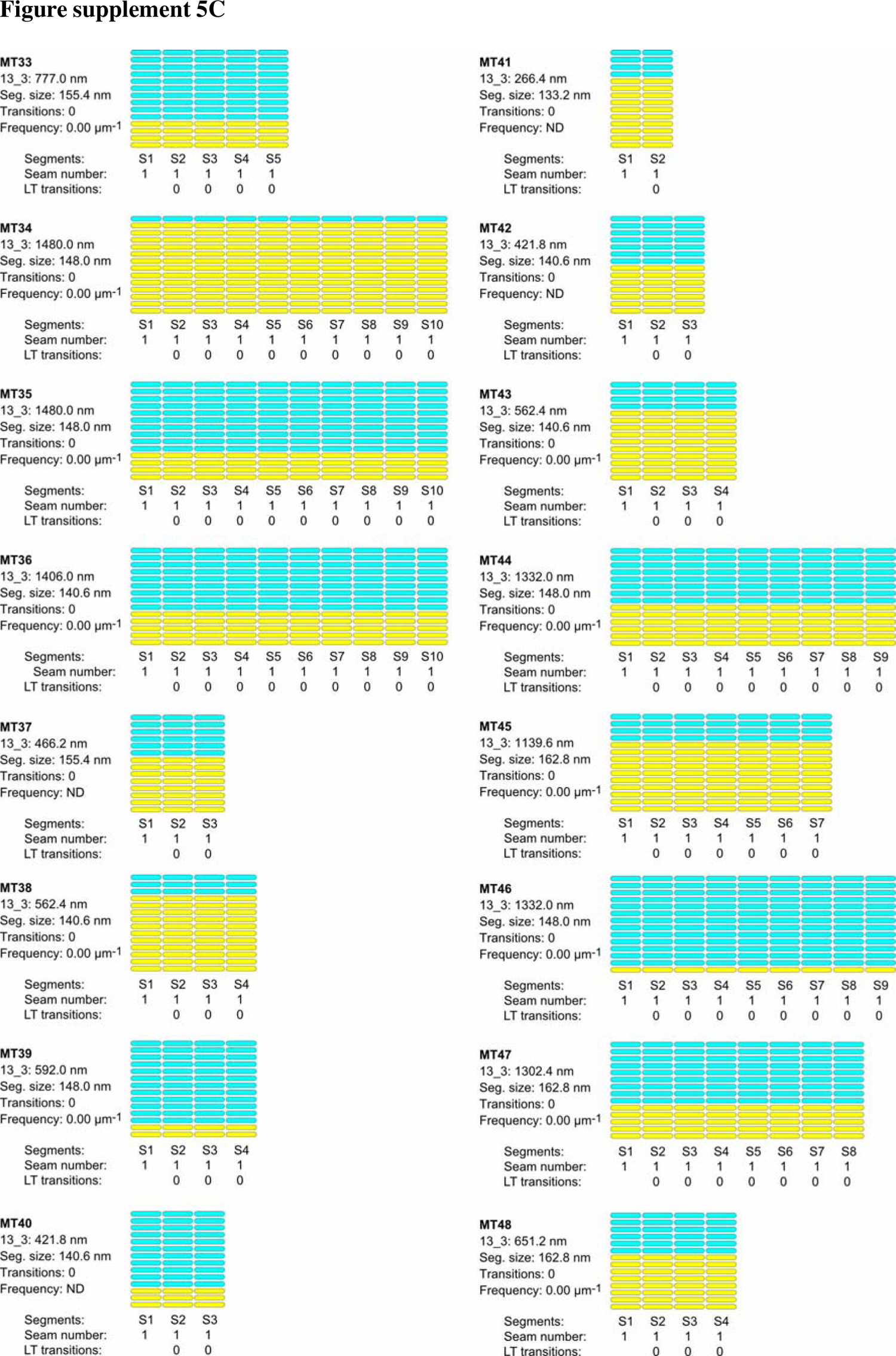
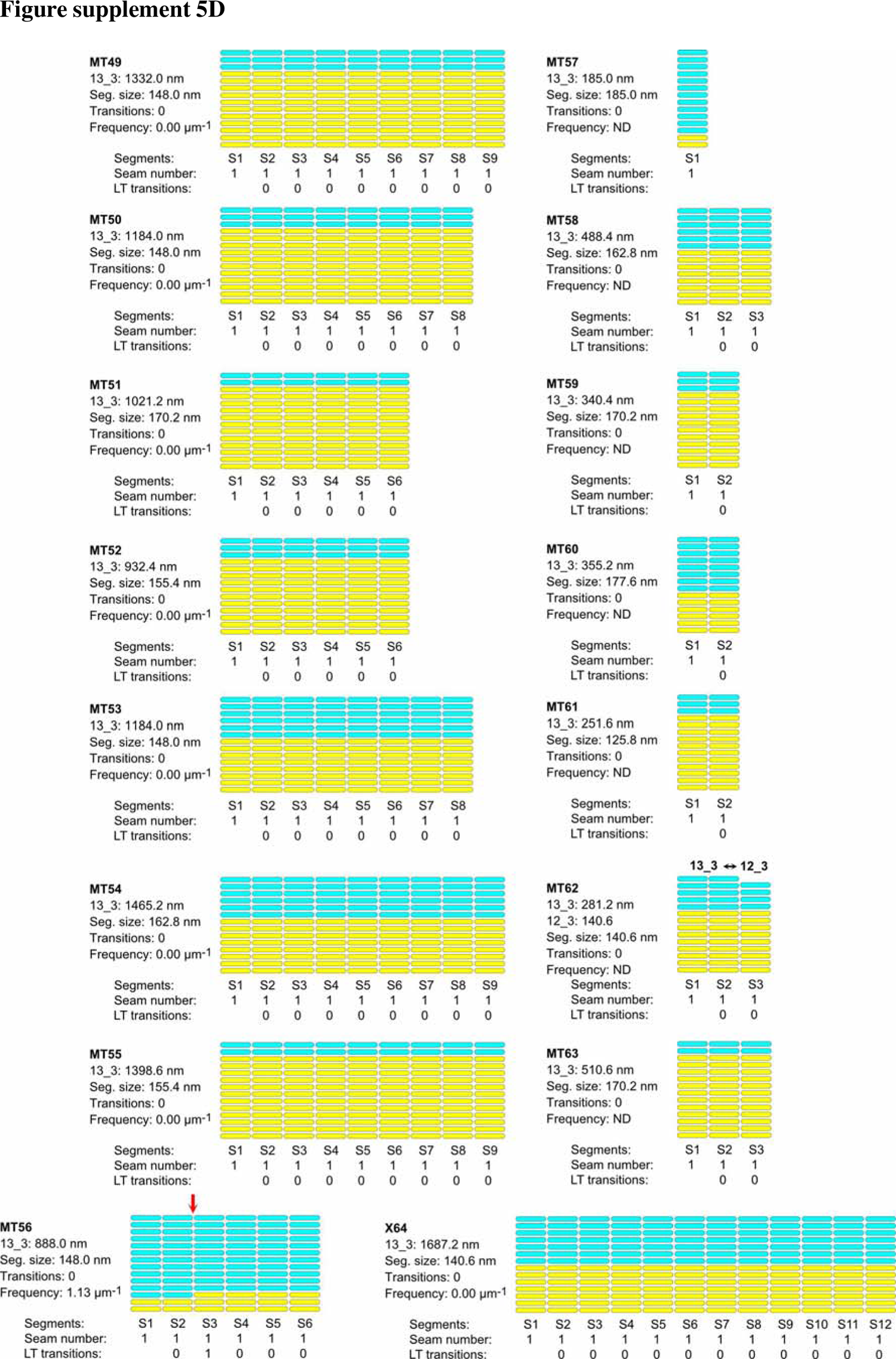
Lattice organization of microtubules assembled in cytoplasmic *Xenopus* egg extracts in the presence of 5% DMSO 64 microtubules (63.5 µm in total length) were analyzed on 5 tomograms acquired on one sample. Number of microtubule segments: 419; number of lateral interactions: 5446 (A type: 414, B type: 5024, ND: 8); lattice type (LT) transitions: 6; protofilament/helical-start (*N/S*) number transitions: 5. Red arrows indicate lattice type transitions in MT2, MT5, MT14, MT18, MT28 and MT56.

**Figure supplement 6.**
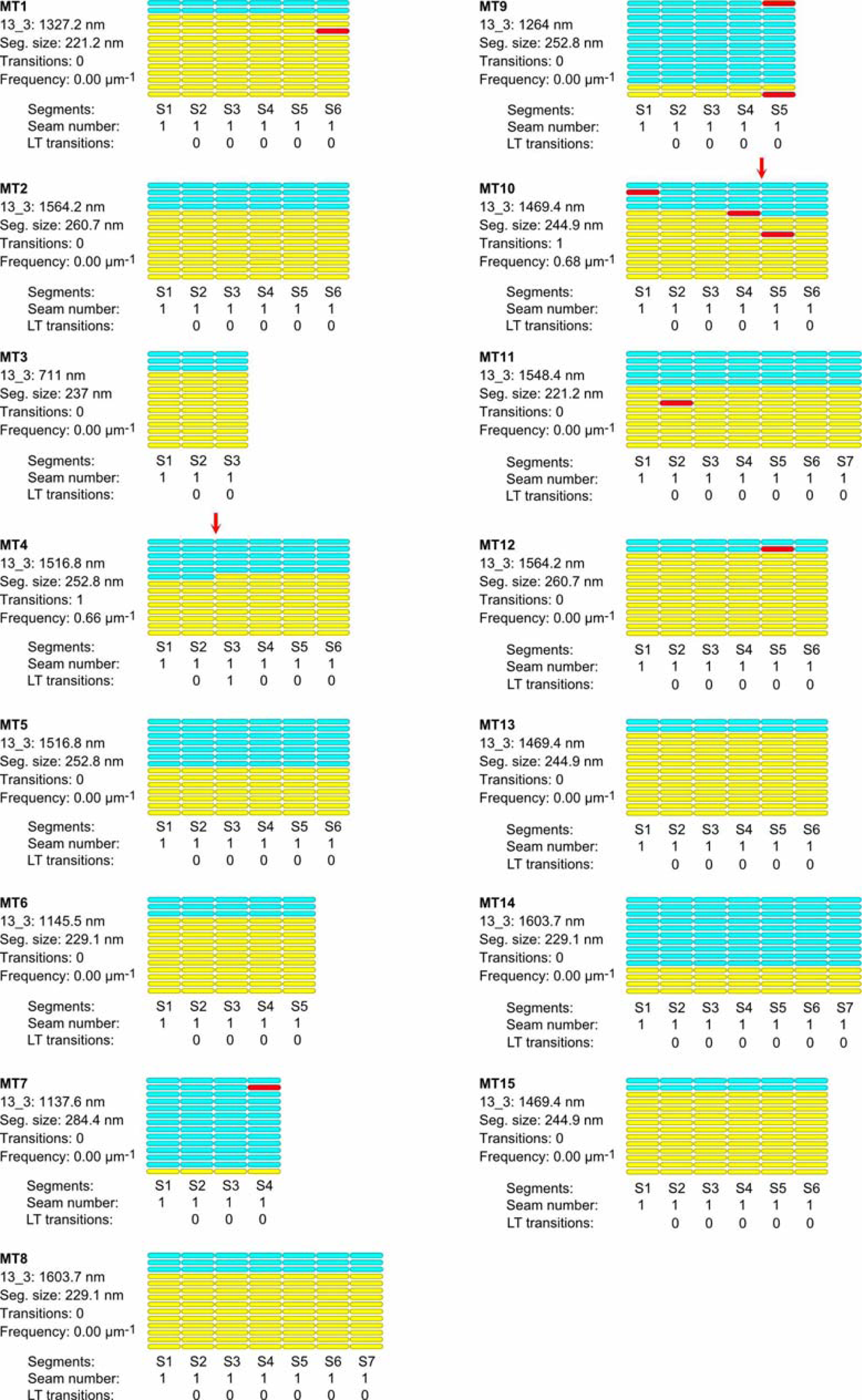
Lattice organization of microtubules assembled in cytoplasmic *Xenopus* egg extracts in the presence of RanQ69L. 15 microtubules (20.6 µm in total length) were analyzed on 1 tomogram acquired on one sample. Number of microtubule segments: 86; number of lateral interactions: 1118 (A type: 8, B type: 1018, ND: 16); lattice type (LT) transitions: 2. Red arrows indicate lattice type transitions in MT4 and MT10.

**Supplementary Table 1.** Characterization of microtubule lattice structure by cryo-electron tomography and segmented sub-tomogram averaging

| Assembly conditions | GTP | GMPCPP | *Xenopus*<br>DMSO | *Xenopus*<br>RanQ69L |
| --- | --- | --- | --- | --- |
| Tomograms | 4 | 6 | 5 | 1 |
| Samples | 2 | 2 | 1 | 1 |
| Microtubules | 24 | 31 | 64 | 15 |
| Total length (μm) | 31.7 | 35.4 | 63.5 | 20.6 |
| Segments | 195 | 239 | 419 | 86 |
| Lateral interactions | 2664 | 3250 | 5446 | 1118 |
| A-type | 461 | 261 | 414 | 8 |
| B-type | 2091 | 2951 | 5024 | 1018 |
| ND | 112 | 38 | 8 | 6 |
| Lattice-type transitions | 119 | 37 | 6 | 2 |
| Frequency (μm<sup>-1</sup>) | 3.69 ± 4.20 | 1.24 ± 1.41 | 0.10 ± 0.28 | 0.13 ± 0.09 |

**Supplementary Table 2.** Protofilament (N) and helix-start number (S)

| N_S | 12_3 | 13_3 | 13_4 | 14_3 | 15_3 | 15_4 |
| --- | --- | --- | --- | --- | --- | --- |
| GTP (%) | 0.99 | 33.07 | - | 64.89 | 1.05 | - |
| GMPCPP (%) | 1.87 | 39.38 | - | 57.06 | - | 1.70 |
| *Xen.* DMSO (%) | 1.16 | 96.30 | 1.59 | 0.95 | - | - |
| *Xen.* RanQ69L (%) | - | 100 | - | - | - | - |

*Xen.*: *Xenopus*

## Supplementary videos

**Figure 2 - Video 1. Dual-axis cryo-electron tomography of microtubules**. Microtubule visualized in cross-section in the single- and dual-axis regions of a cryo-electron tomogram. Related to Figure 2 - figure supplement 1C.

**Figure 2 - Video 2. Sub-tomogram average of a 14_3 mono-seam microtubule**. Yellow spheres are placed on kinesin-motor domain densities and cyan spheres in between along one turn of the 3-start helix.

**Figure 3 - Video 3. SSTA of a 13_3 multi-seam microtubule**. Sub-tomogram average of the 13_3 microtubule in Figure 3A (full volume), followed by SSTA of the microtubule in 4 segments of 331.8 nm in length (Figure 3B). Red spheres are placed onto aberrant protofilaments.

**Figure 7 - Video 4. Cytoplasmic extract microtubules decorated with kinesin-motor domains.** Slices through a cryo-electron tomogram of *Xenopus* egg cytoplasm showing microtubules decorated with kinesin-motor domains.

**Figure 8 - Video 5. SSTA of a 13_3 microtubule assembled in a cytoplasmic extract**. SSTA of the 13_3 microtubule in Figure 8A in 310.8 nm long segments (S3-S5, Figure 8A, *top*), and 148.0 nm long segments (S7-S10, Figure 8A, *bottom*).

**Figure 9 - Video 6. Sub-tomogram averages of microtubules with different protofilament and/or helix-start numbers.** Sub-tomogram averages of the 12_2, 12_3, 13_4, 14_3 microtubules in Figure 9. Arrows point to local dislocations in the 12_2 and 13_4 microtubules.

**Figure 10 - Video 7. Microtubule growth-induced mechanism of lattice heterogeneity**. This video depicts how holes of an odd or even number of subunits arise during microtubule polymerization, with the formation of lateral interactions without longitudinal ones at the tip of the growing microtubule.

## Notes

### Competing Interest Statement

The authors have declared no competing interest.

### Summary of Updates

Additional data have been added on microtubules polymerized from purified tubulin in the presence of GMPCPP. Holes are visualized in raw tomograms (new Figure 5) and artefacts that limit their visualization are presented in new Figure 6. Data on microtubules polymerized in Xenopus egg cytoplasmic extract assembled in the presence of RanQ69L are presented as a control to DMSO-induced asters (new Supplementary Figure 6).

## References

1. Akhmanova, A., & Steinmetz, M. O. (2008). Tracking the ends: A dynamic protein network controls the fate of microtubule tips. Nature Reviews. Molecular Cell Biology, 9(4), 309–322. https://doi.org/10.1038/nrm2369

2. Amos, L. A., & Klug, A. (1974). Arrangement of subunits in flagellar microtubules. Journal of Cell Science, 14(3), 523–549 https://doi.org/10.1242/jcs.14.3.523

3. Ashford, A. J., & Hyman, A. A. (2006). Chapter 22—Preparation of Tubulin from Porcine Brain. In J. E. Celis (Éd.), Cell Biology (Third Edition) (p. 155–160). Academic Press. https://doi.org/10.1016/B978-012164730-8/50094-0

4. Böhler, A., Vermeulen, B. J. A., Würtz, M., Zupa, E., Pfeffer, S., & Schiebel, E. (2021). The gamma-tubulin ring complex: Deciphering the molecular organization and assembly mechanism of a major vertebrate microtubule nucleator. BioEssays, 43(8), 2100114. https://doi.org/10.1002/bies.202100114

5. Bowne-Anderson, H., Hibbel, A., & Howard, J. (2015). Regulation of Microtubule Growth and Catastrophe: Unifying Theory and Experiment. Trends in cell biology, 25(12), 769–779. https://doi.org/10.1016/j.tcb.2015.08.009

6. Bowne-Anderson, H., Zanic, M., Kauer, M., & Howard, J. (2013). Microtubule dynamic instability: A new model with coupled GTP hydrolysis and multistep catastrophe. *BioEssays: News and Reviews in Molecular*, Cellular and Developmental Biology, 35(5), 452–461. https://doi.org/10.1002/bies.201200131

7. Brouhard, G. J. (2015). Dynamic instability 30 years later: Complexities in microtubule growth and catastrophe. Molecular Biology of the Cell, 26(7), 1207–1210. https://doi.org/10.1091/mbc.E13-10-0594

8. Brouhard, G. J., Stear, J. H., Noetzel, T. L., Al-Bassam, J., Kinoshita, K., Harrison, S. C., Howard, J., & Hyman, A. A. (2008). XMAP215 is a processive microtubule polymerase. Cell, 132(1), 79–88. https://doi.org/10.1016/j.cell.2007.11.043

9. Carazo-Salas, R. E., Guarguaglini, G., Gruss, O. J., Segref, A., Karsenti, E., & Mattaj, I. W. (1999). Generation of GTP-bound Ran by RCC1 is required for chromatin-induced mitotic spindle formation. Nature, 400(6740), 178–181. https://doi.org/10.1038/22133

10. Castoldi, M., & Popov, A. V. (2003). Purification of brain tubulin through two cycles of polymerization–depolymerization in a high-molarity buffer. Protein Expression and Purification, 32(1), 83–88. https://doi.org/10.1016/S1046-5928(03)00218-3

11. Chaaban, S., & Brouhard, G. J. (2017). A microtubule bestiary: Structural diversity in tubulin polymers. Molecular Biology of the Cell, 28(22), 2924–2931. https://doi.org/10.1091/mbc.E16-05-0271

12. Chen, J., Noble, A. J., Kang, J. Y., & Darst, S. A. (2019). Eliminating effects of particle adsorption to the air/water interface in single-particle cryo-electron microscopy: Bacterial RNA polymerase and CHAPSO. Journal of Structural Biology: X, 1, 100005. https://doi.org/10.1016/j.yjsbx.2019.100005

13. Chen, S., Li, J., Vinothkumar, K. R., & Henderson, R. (2022). Interaction of human erythrocyte catalase with air–water interface in cryoEM. Microscopy, 71(Supplement_1), i51–i59. https://doi.org/10.1093/jmicro/dfab037

14. Chrétien, D., & Fuller, S. D. (2000). Microtubules switch occasionally into unfavorable configurations during elongation. Journal of Molecular Biology, 298(4), 663–676. https://doi.org/10.1006/jmbi.2000.3696

15. Chrétien, D., Metoz, F., Verde, F., Karsenti, E., & Wade, R. (1992). Lattice defects in microtubules: Protofilament numbers vary within individual microtubules. Journal of Cell Biology, 117(5), 1031–1040. https://doi.org/10.1083/jcb.117.5.1031

16. Chrétien, D., & Wade, R. H. (1991). New data on the microtubule surface lattice. Biology of the Cell, 71(1-2), 161–174. https://doi.org/10.1016/0248-4900(91)90062-r

17. Cleary, J. M., & Hancock, W. O. (2021). Molecular mechanisms underlying microtubule growth dynamics. Current Biology: CB, 31(10), R560–R573. https://doi.org/10.1016/j.cub.2021.02.035

18. Crepeau, R. H., McEwen, B., & Edelstein, S. J. (1978). Differences in alpha and beta polypeptide chains of tubulin resolved by electron microscopy with image reconstruction. Proceedings of the National Academy of Sciences of the United States of America, 75(10), 5006–5010. https://doi.org/10.1073/pnas.75.10.5006

19. Debs, G. E., Cha, M., Liu, X., Huehn, A. R., & Sindelar, C. V. (2020). Dynamic and asymmetric fluctuations in the microtubule wall captured by high-resolution cryoelectron microscopy. Proceedings of the National Academy of Sciences of the United States of America, 117(29), 16976–16984. https://doi.org/10.1073/pnas.2001546117

20. des Georges, A., Katsuki, M., Drummond, D. R., Osei, M., Cross, R. A., & Amos, L. A. (2008). Mal3, the Schizosaccharomyces pombe homolog of EB1, changes the microtubule lattice. Nature Structural & Molecular Biology, 15(10), 1102–1108. https://doi.org/10.1038/nsmb.1482

21. Dimitrov, A., Quesnoit, M., Moutel, S., Cantaloube, I., Poüs, C., & Perez, F. (2008). Detection of GTP-tubulin conformation in vivo reveals a role for GTP remnants in microtubule rescues. *Science (New York*, N.Y.), 322(5906), 1353–1356. https://doi.org/10.1126/science.1165401

22. D’Imprima, E., Floris, D., Joppe, M., Sánchez, R., Grininger, M., & Kühlbrandt, W. (2019). Protein denaturation at the air-water interface and how to prevent it. eLife, 8, e42747. https://doi.org/10.7554/eLife.42747

23. Duchesne, L., Gentili, D., Comes-Franchini, M., & Fernig, D. G. (2008). Robust Ligand Shells for Biological Applications of Gold Nanoparticles. Langmuir, 24(23), 13572–13580. https://doi.org/10.1021/la802876u

24. Edelstein, A. D., Tsuchida, M. A., Amodaj, N., Pinkard, H., Vale, R. D., & Stuurman, N. (2014). Advanced methods of microscope control using μManager software. Journal of Biological Methods, 1(2), e10–e10. https://doi.org/10.14440/jbm.2014.36

25. Farmer, V., Arpağ, G., Hall, S. L., & Zanic, M. (2021). XMAP215 promotes microtubule catastrophe by disrupting the growing microtubule end. Journal of Cell Biology, 220(10), e202012144. https://doi.org/10.1083/jcb.202012144

26. Gibeaux, R., & Heald, R. (2019). The Use of Cell-Free Xenopus Extracts to Investigate Cytoplasmic Events. Cold Spring Harbor Protocols, 2019(6). https://doi.org/10.1101/pdb.top097048

27. Good, M. C., & Heald, R. (2018). Preparation of Cellular Extracts from Xenopus Eggs and Embryos. Cold Spring Harbor Protocols, 2018(6), pdb.prot097055. https://doi.org/10.1101/pdb.prot097055

28. Guesdon, A., Bazile, F., Buey, R. M., Mohan, R., Monier, S., García, R. R., Angevin, M., Heichette, C., Wieneke, R., Tampé, R., Duchesne, L., Akhmanova, A., Steinmetz, M. O., & Chrétien, D. (2016). EB1 interacts with outwardly curved and straight regions of the microtubule lattice. Nature Cell Biology, 18(10), 1102–1108. https://doi.org/10.1038/ncb3412

29. Guesdon, A., Blestel, S., Kervrann, C., & Chrétien, D. (2013). Single versus dual-axis cryo-electron tomography of microtubules assembled in vitro: Limits and perspectives. Journal of Structural Biology, 181(2), 169–178. https://doi.org/10.1016/j.jsb.2012.11.004

30. Hagen, W. J. H., Wan, W., & Briggs, J. A. G. (2017). Implementation of a cryo-electron tomography tilt-scheme optimized for high resolution subtomogram averaging. Journal of Structural Biology, 197(2), 191–198. https://doi.org/10.1016/j.jsb.2016.06.007

31. Helmke, K. J., & Heald, R. (2014). TPX2 levels modulate meiotic spindle size and architecture in Xenopus egg extracts. The Journal of Cell Biology, 206(3), 385–393. https://doi.org/10.1083/jcb.201401014

32. Howes, S. C., Geyer, E. A., LaFrance, B., Zhang, R., Kellogg, E. H., Westermann, S., Rice, L. M., & Nogales, E. (2017). Structural differences between yeast and mammalian microtubules revealed by cryo-EM. The Journal of Cell Biology, 216(9), 2669–2677. https://doi.org/10.1083/jcb.201612195

33. Hunyadi, V., Chrétien, D., & Jánosi, I. M. (2005). Mechanical stress induced mechanism of microtubule catastrophes. Journal of Molecular Biology, 348(4), 927–938. https://doi.org/10.1016/j.jmb.2005.03.019

34. Kikkawa, M., Ishikawa, T., Nakata, T., Wakabayashi, T., & Hirokawa, N. (1994). Direct visualization of the microtubule lattice seam both in vitro and in vivo. The Journal of Cell Biology, *127*(6 Pt 2), 1965-1971. https://doi.org/10.1083/jcb.127.6.1965

35. Klebl, D. P., Wang, Y., Sobott, F., Thompson, R. F., & Muench, S. P. (2022). It started with a Cys: Spontaneous cysteine modification during cryo-EM grid preparation. Frontiers in Molecular Biosciences, 9. https://www.frontiersin.org/articles/10.3389/fmolb.2022.945772

36. Kremer, J. R., Mastronarde, D. N., & McIntosh, J. R. (1996). Computer Visualization of Three-Dimensional Image Data Using IMOD. Journal of Structural Biology, 116(1), 71–76. https://doi.org/10.1006/jsbi.1996.0013

37. Li, B., Zhu, D., Shi, H., & Zhang, X. (2021). Effect of charge on protein preferred orientation at the air–water interface in cryo-electron microscopy. Journal of Structural Biology, 213(4), 107783. https://doi.org/10.1016/j.jsb.2021.107783

38. Manka, S. W., & Moores, C. A. (2018). Microtubule structure by cryo-EM: Snapshots of dynamic instability. Essays in Biochemistry, 62(6), 737–751. https://doi.org/10.1042/EBC20180031

39. Mastronarde, D. N. (1997). Dual-Axis Tomography: An Approach with Alignment Methods That Preserve Resolution. Journal of Structural Biology, 120(3), 343–352. https://doi.org/10.1006/jsbi.1997.3919

40. Maurer, S. P., Fourniol, F. J., Bohner, G., Moores, C. A., & Surrey, T. (2012). EBs recognize a nucleotide-dependent structural cap at growing microtubule ends. Cell, 149(2), 371–382. https://doi.org/10.1016/j.cell.2012.02.049

41. McEwen, B., & Edelstein, S. J. (1980). Evidence for a mixed lattice in microtubules reassembled in vitro. Journal of Molecular Biology, 139(2), 123–143. https://doi.org/10.1016/0022-2836(80)90300-9

42. McIntosh, J. R., Morphew, M. K., Grissom, P. M., Gilbert, S. P., & Hoenger, A. (2009). Lattice structure of cytoplasmic microtubules in a cultured Mammalian cell. Journal of Molecular Biology, 394(2), 177–182. https://doi.org/10.1016/j.jmb.2009.09.033

43. McIntosh, J. R., O’Toole, E., Morgan, G., Austin, J., Ulyanov, E., Ataullakhanov, F., & Gudimchuk, N. (2018). Microtubules grow by the addition of bent guanosine triphosphate tubulin to the tips of curved protofilaments. Journal of Cell Biology, 217(8), 2691–2708. https://doi.org/10.1083/jcb.201802138

44. Mitchison, T., & Kirschner, M. (1984). Dynamic instability of microtubule growth. Nature, 312(5991), 237–242. https://doi.org/10.1038/312237a0

45. Murray, A. W. (1991). Cell cycle extracts. Methods in Cell Biology, 36, 581–605.

46. Nicastro, D., Schwartz, C., Pierson, J., Gaudette, R., Porter, M. E., & McIntosh, J. R. (2006). The Molecular Architecture of Axonemes Revealed by Cryoelectron Tomography. Science, 313(5789), 944–948. https://doi.org/10.1126/science.1128618

47. Olieric, N., Kuchen, M., Wagen, S., Sauter, M., Crone, S., Edmondson, S., Frey, D., Ostermeier, C., Steinmetz, M. O., & Jaussi, R. (2010). Automated seamless DNA co-transformation cloning with direct expression vectors applying positive or negative insert selection. BMC Biotechnology, 10(1), 56. https://doi.org/10.1186/1472-6750-10-56

48. Pantaloni, D., & Carlier, M.-F. (1986). Involvement of Guanosine Triphosphate (GTP) Hydrolysis in the Mechanism of Tubulin Polymerization: Regulation of Microtubule Dynamics at Steady State by a GTP Cap. Annals of the New York Academy of Sciences, 466(1), 496–509. https://doi.org/10.1111/j.1749-6632.1986.tb38427.x

49. Rai, A., Liu, T., Katrukha, E. A., Estévez-Gallego, J., Manka, S. W., Paterson, I., Díaz, J. F., Kapitein, L. C., Moores, C. A., & Akhmanova, A. (2021). Lattice defects induced by microtubule-stabilizing agents exert a long-range effect on microtubule growth by promoting catastrophes. Proceedings of the National Academy of Sciences, 118(51), e2112261118. https://doi.org/10.1073/pnas.2112261118

50. Rothwell, S. W., Grasser, W. A., & Murphy, D. B. (1986). End-to-end annealing of microtubules in vitro. The Journal of Cell Biology, 102(2), 619–627. https://doi.org/10.1083/jcb.102.2.619

51. Sawin, K. E., & Mitchison, T. J. (1994). Microtubule flux in mitosis is independent of chromosomes, centrosomes, and antiparallel microtubules. Molecular Biology of the Cell, 5(2), 217–226. https://doi.org/10.1091/mbc.5.2.217

52. Schaedel, L., Triclin, S., Chrétien, D., Abrieu, A., Aumeier, C., Gaillard, J., Blanchoin, L., Théry, M., & John, K. (2019). Lattice defects induce microtubule self-renewal. Nature Physics, 15(8), 830–838. https://doi.org/10.1038/s41567-019-0542-4

53. Song, Y. H., & Mandelkow, E. (1993). Recombinant kinesin motor domain binds to beta-tubulin and decorates microtubules with a B surface lattice. Proceedings of the National Academy of Sciences of the United States of America, 90(5), 1671–1675. https://doi.org/10.1073/pnas.90.5.1671

54. Song, Y. H., & Mandelkow, E. (1995). The anatomy of flagellar microtubules: Polarity, seam, junctions, and lattice. The Journal of Cell Biology, 128(1-2), 81[94. https://doi.org/10.1083/jcb.128.1.81

55. Sosa, H., Hoenger, A., & Milligan, R. A. (1997). Three different approaches for calculating the three-dimensional structure of microtubules decorated with kinesin motor domains. Journal of Structural Biology, 118(2), 149–158. https://doi.org/10.1006/jsbi.1997.3851

56. Théry, M., & Blanchoin, L. (2021). Microtubule self-repair. Current Opinion in Cell Biology, 68, 144–154. https://doi.org/10.1016/j.ceb.2020.10.012

57. Vitre, B., Coquelle, F. M., Heichette, C., Garnier, C., Chrétien, D., & Arnal, I. (2008). EB1 regulates microtubule dynamics and tubulin sheet closure in vitro. Nature Cell Biology, 10(4), 415–421. https://doi.org/10.1038/ncb1703

58. Wade, R. H., & Chrétien, D. (1993). Cryoelectron microscopy of microtubules. Journal of Structural Biology, 110(1), 1–27. https://doi.org/10.1006/jsbi.1993.1001

59. Zabeo, D., Heumann, J. M., Schwartz, C. L., Suzuki-Shinjo, A., Morgan, G., Widlund, P. O., & Höög, J. L. (2018). A lumenal interrupted helix in human sperm tail microtubules. Scientific Reports, 8(1), 2727. https://doi.org/10.1038/s41598-018-21165-8

60. Zhang, R., Alushin, G. M., Brown, A., & Nogales, E. (2015). Mechanistic Origin of Microtubule Dynamic Instability and Its Modulation by EB Proteins. Cell, 162(4), 849–859. https://doi.org/10.1016/j.cell.2015.07.012

