## Supplementary figures and images for "Changes in seam number and location induce holes within microtubules assembled from porcine brain tubulin and in *Xenopus* egg cytoplasmic extracts"

### Figure supplement 1

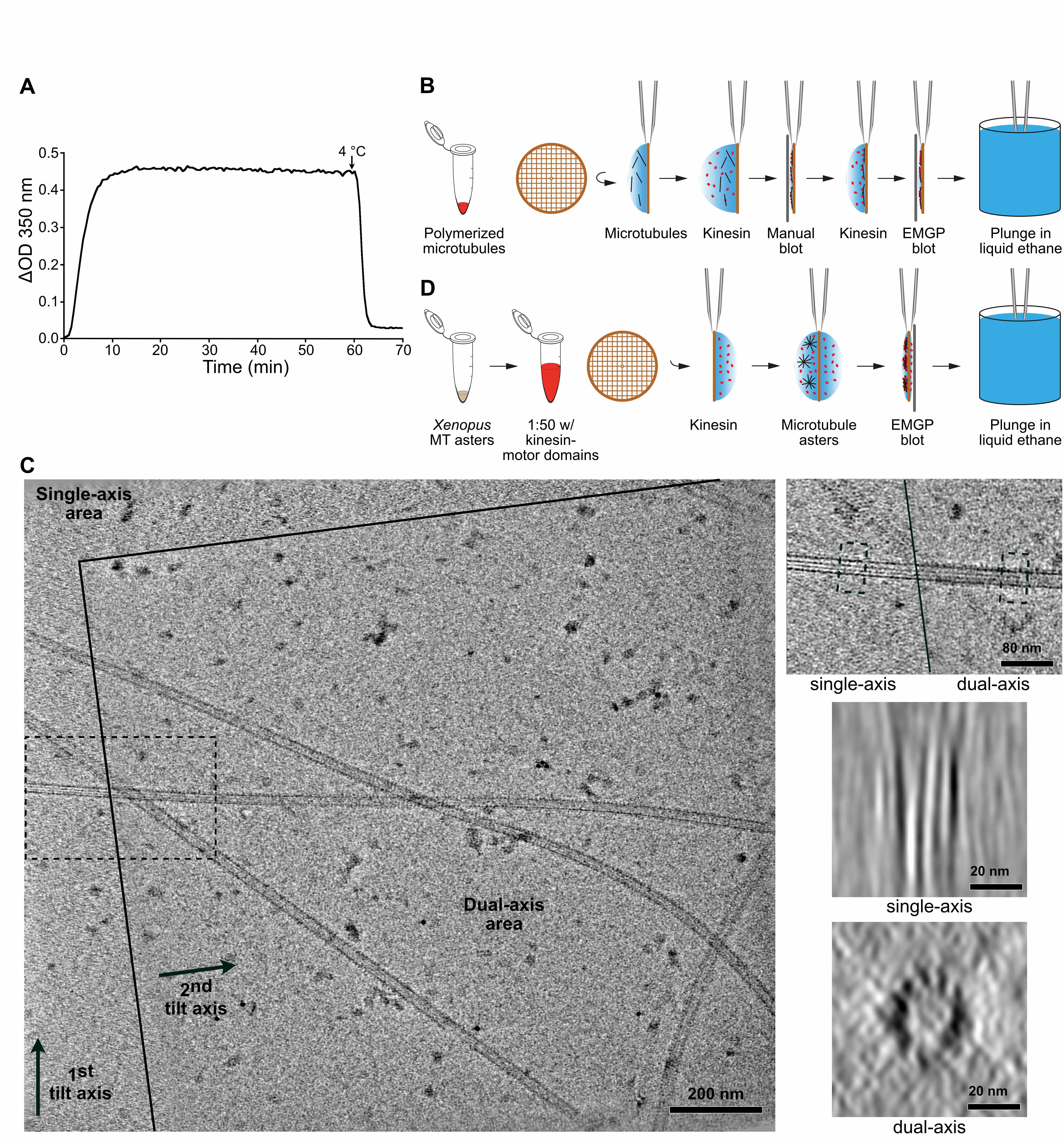

### Figure supplement 2

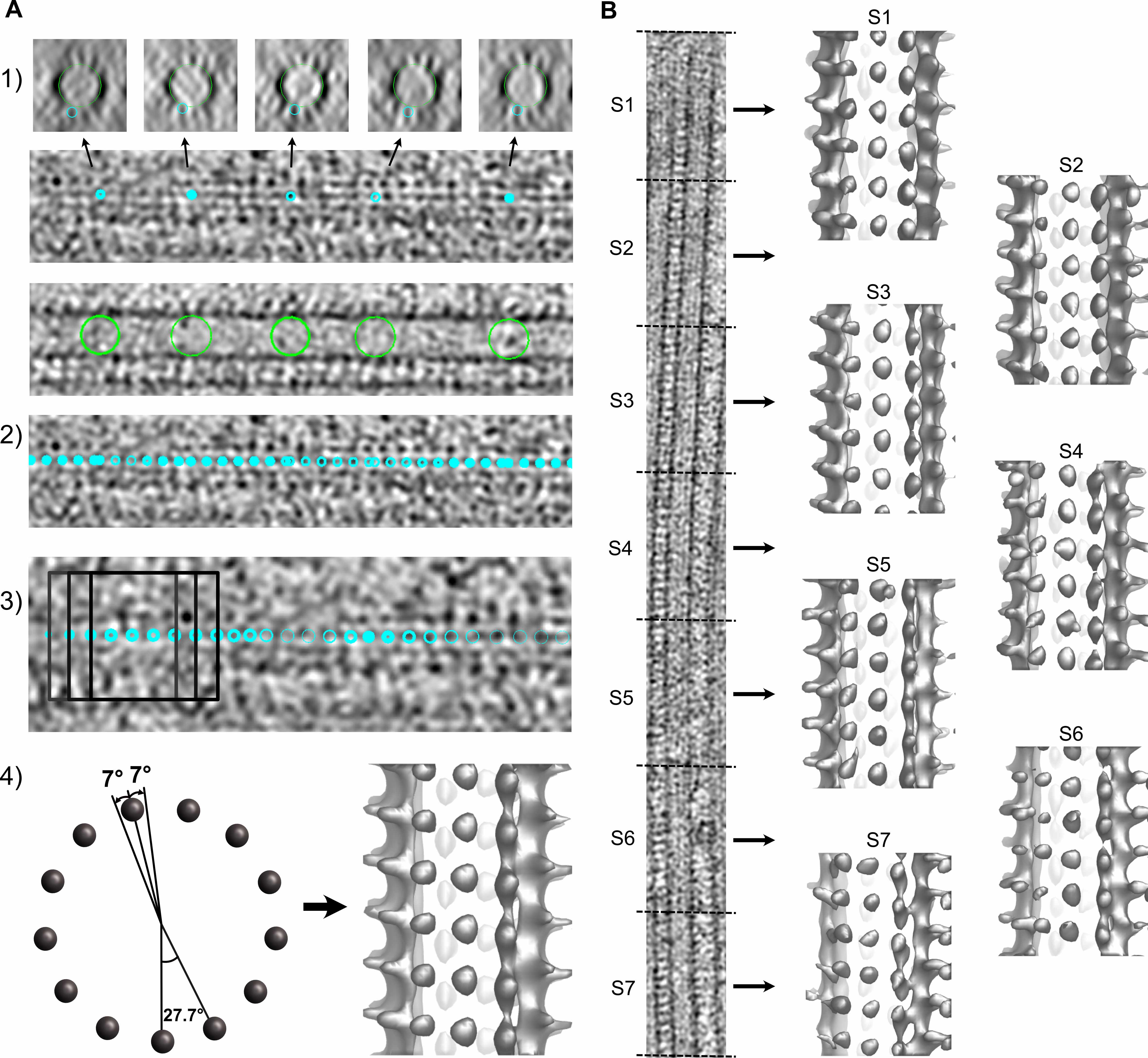

### Figure supplement 3A

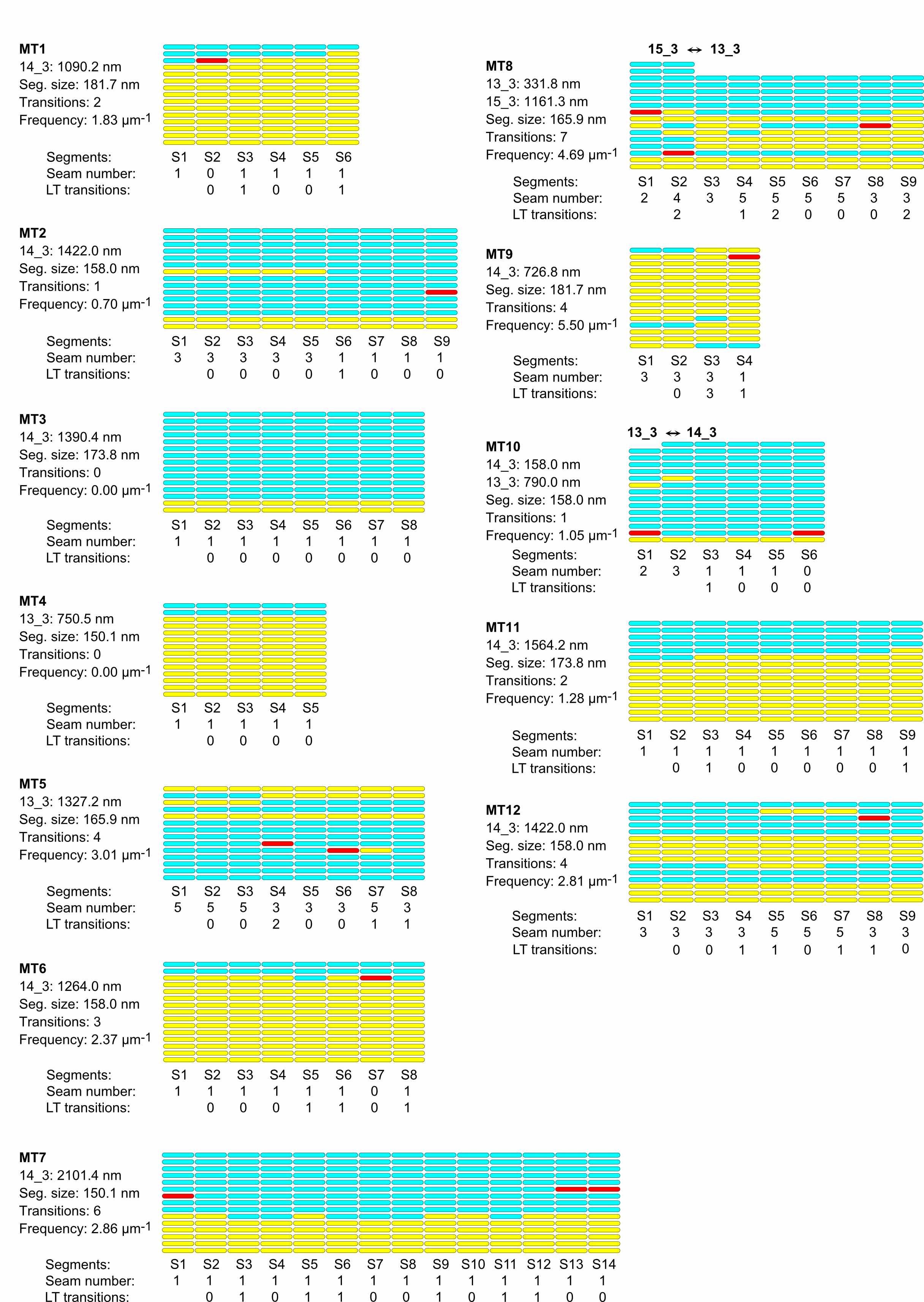

### Figure supplement 3B

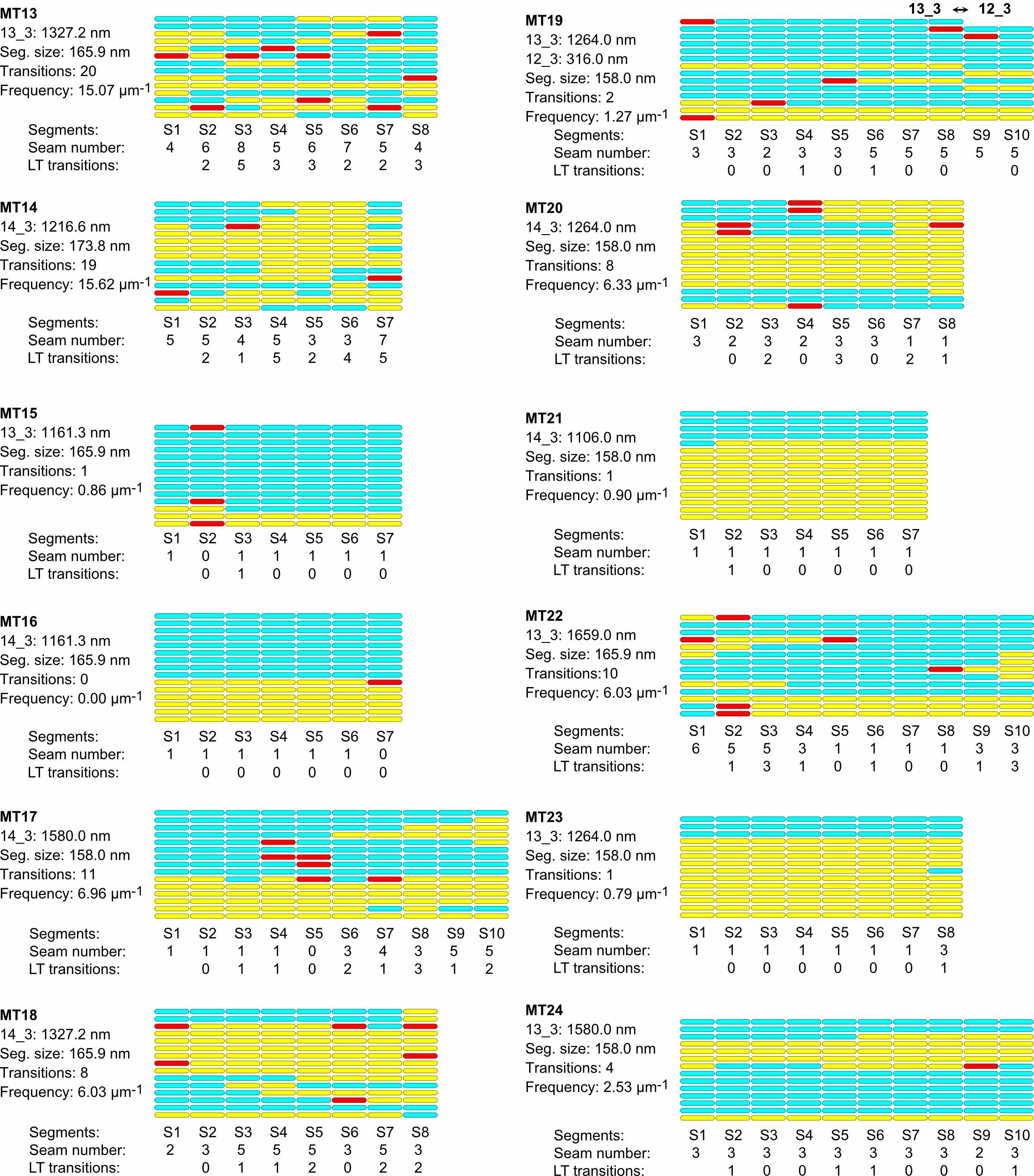

### Figure supplement 4A

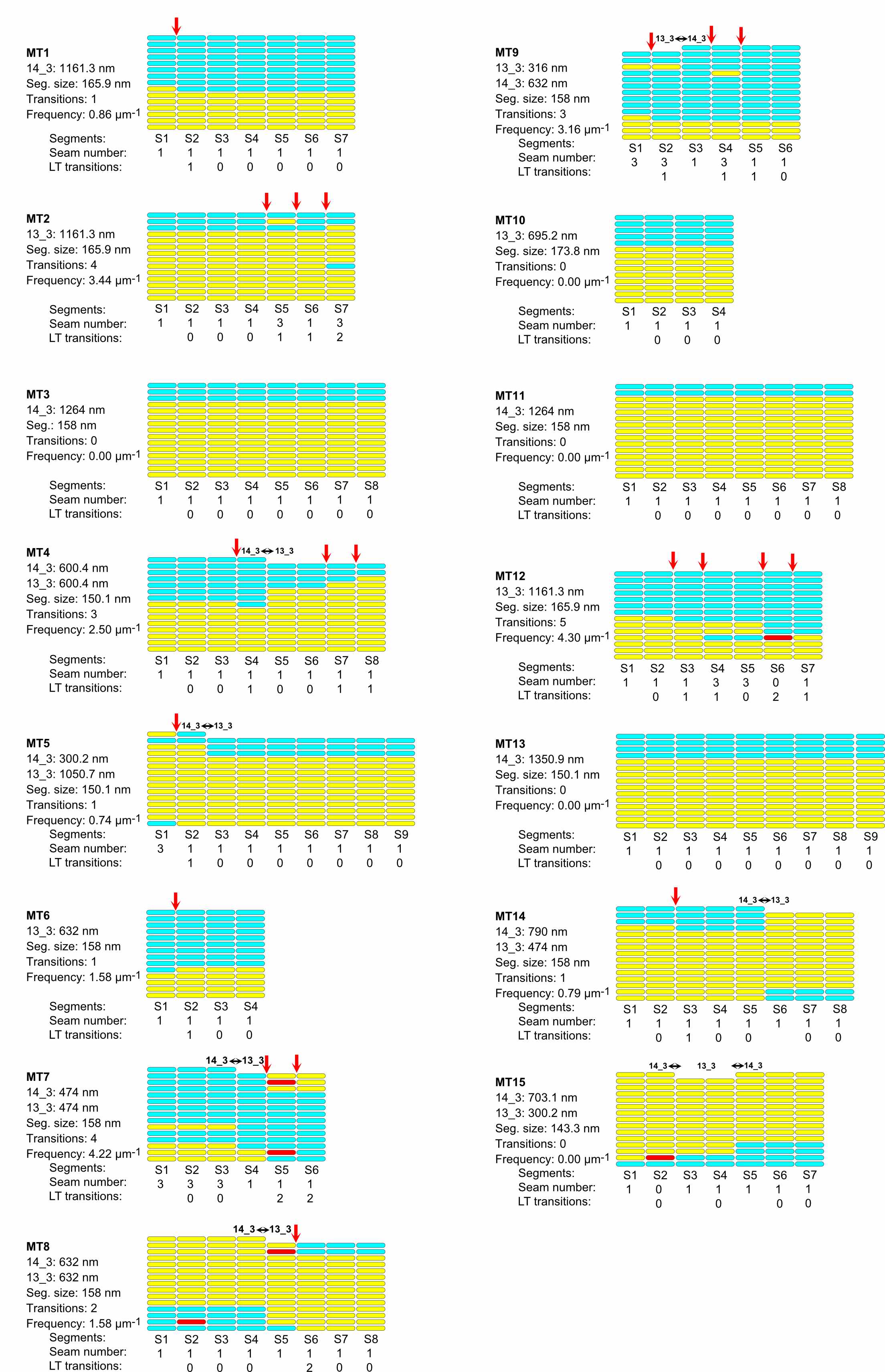

### Figure supplement 4B

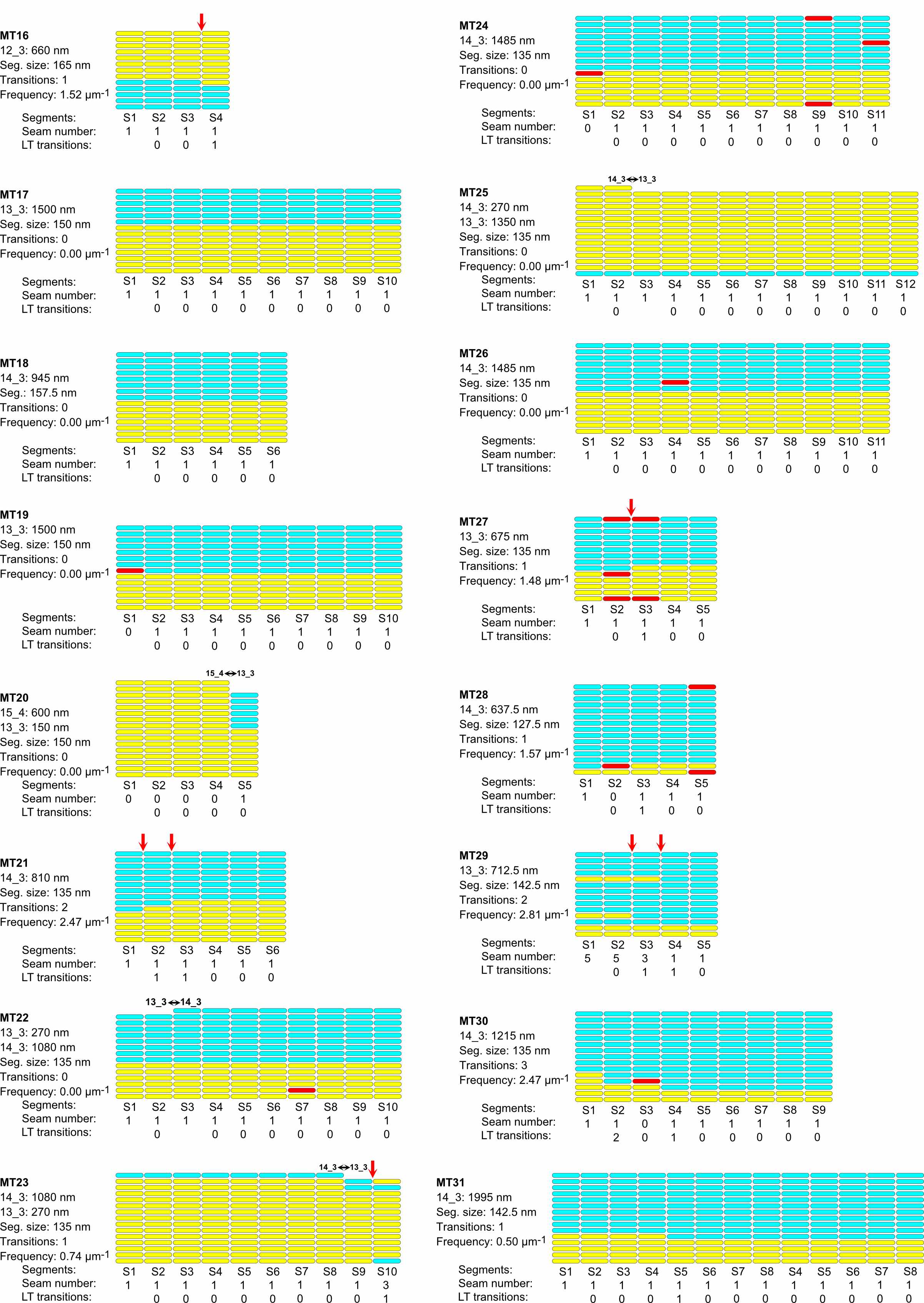

### Figure supplement 5A

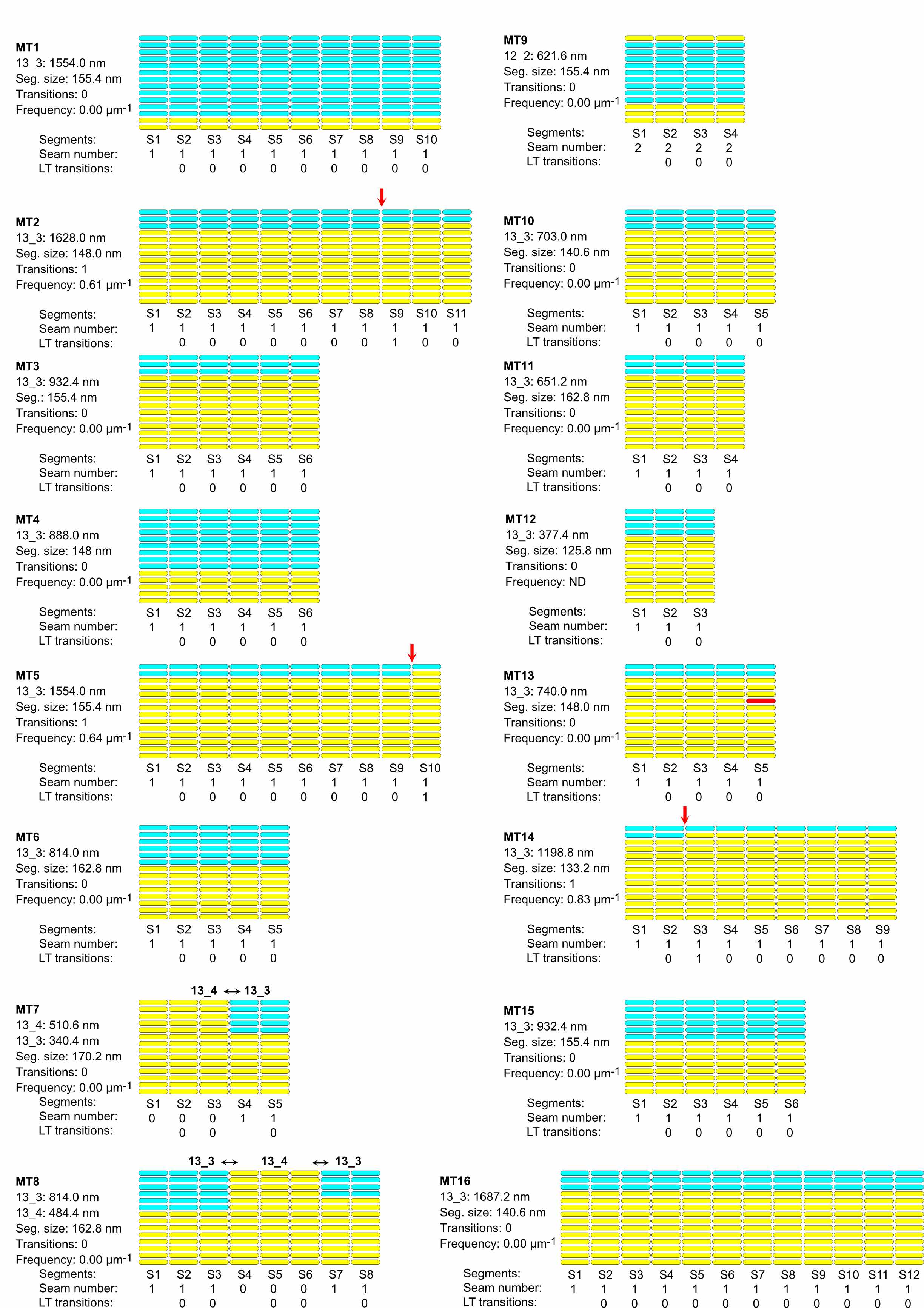

### Figure supplement 5B

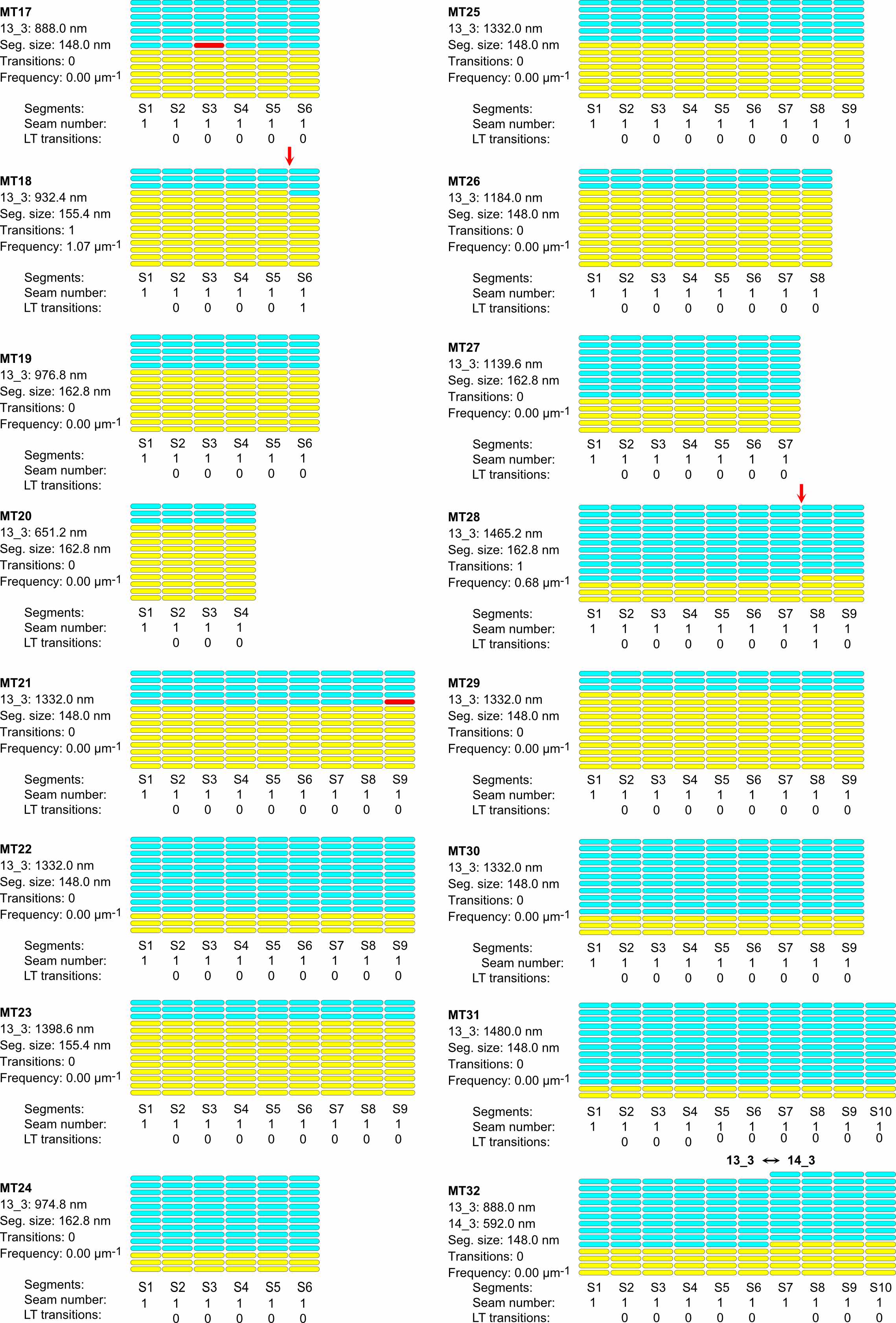

### Figure supplement 5C

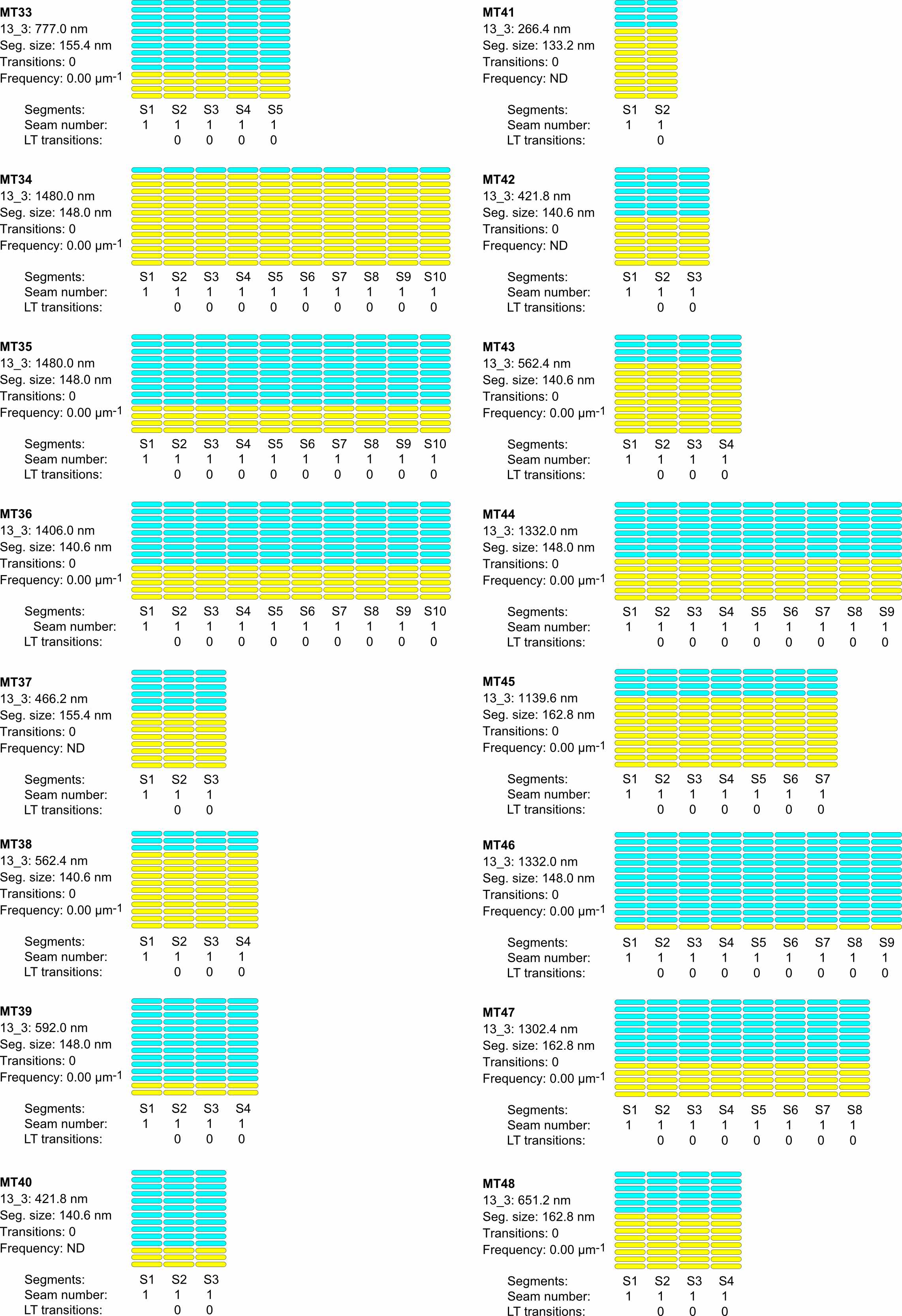

### Figure supplement 5D

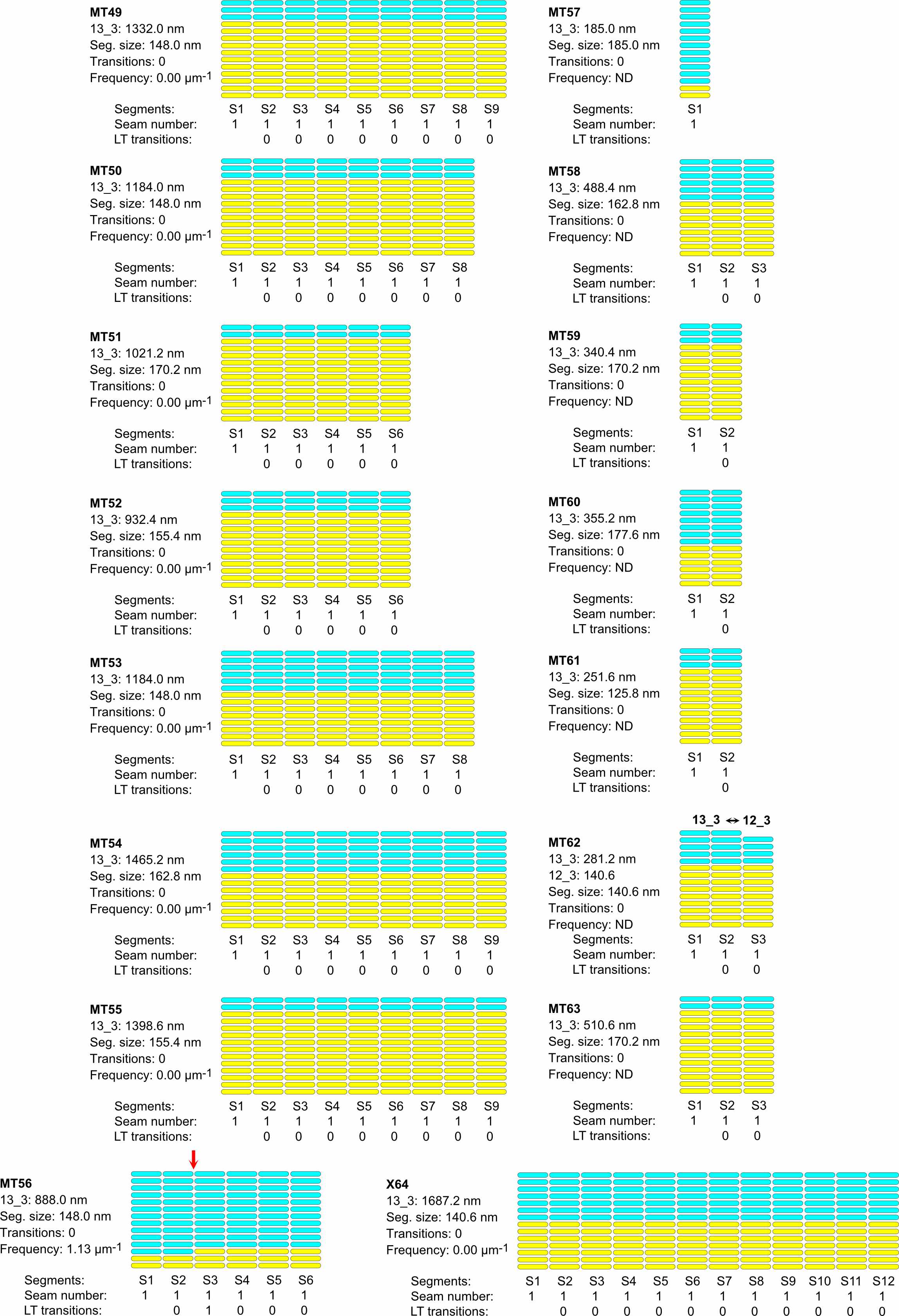

### Figure supplement 6

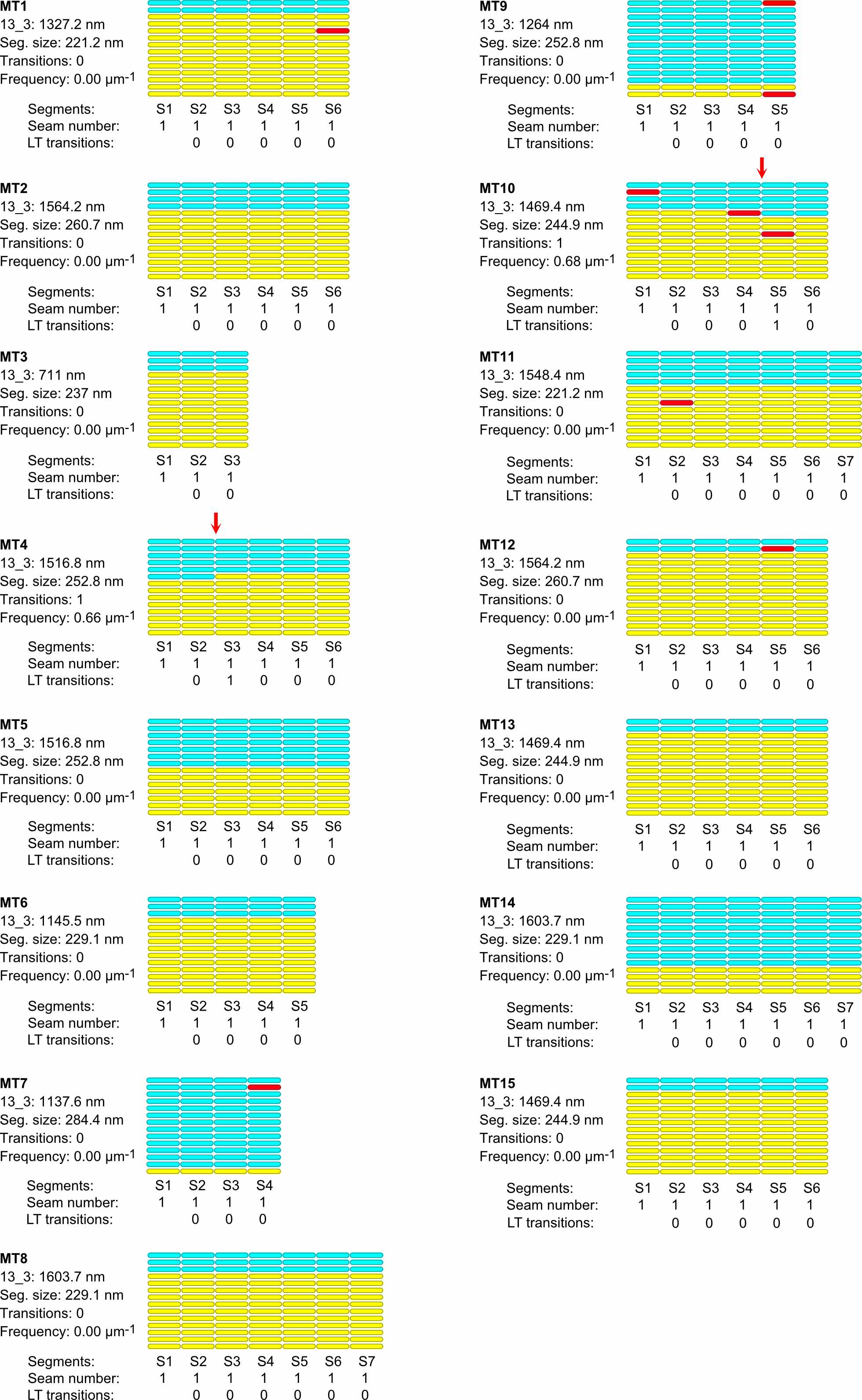
