## Supplementray Table 1 for "Changes in seam number and location induce holes within microtubules assembled from porcine brain tubulin and in *Xenopus* egg cytoplasmic extracts"

**Supplementary Table 1. Characterization of microtubule lattice structure by cryo-electron tomography and segmented sub-tomogram averaging**

| Assembly conditions | GTP | GMPCPP | *Xenopus* DMSO | *Xenopus* Ran |
| --- | --- | --- | --- | --- |
| Tomograms | 4 |  | 5 | 1 |
| Samples | 2 | 2 | 1 | 1 |
| Microtubules | 24 |  | 64 | 15 |
| Total length (µm) | 31.7 |  | 63.5 | 20.6 |
| Segments | 195 |  | 419 | 86 |
| Lateral interactions | 2664 |  | 5446 | 1118 |
| A-type | 461 |  | 414 | 8 |
| B-type | 2091 |  | 5024 | 1018 |
| ND | 112 |  | 8 | 6 |
| Lattice-type transitions | 119 |  | 6 | 2 |
| Frequency (µm^-1^) | 3.69 ± 4.20 |  | 0.10 ± 0.28 | 0.13 ± 0.09 |

**Supplementary Table 2. Protofilament (N) and helix-start number (S)**

| N_S | 12_2 | 12_3 | 13_3 | 13_4 | 14_3 | 14_4 | 15_3 | 15_4 |
| --- | --- | --- | --- | --- | --- | --- | --- | --- |
| GTP (%) |  |  |  |  |  |  |  |  |
| GMPCPP (%) |  |  |  |  |  |  |  |  |
| *Xen.* DMSO (%) |  |  |  |  |  |  |  |  |
| *Xen.* Ran (%) |  |  |  |  |  |  |  |  |
